# Somatic genetics analysis of sleep in adult mice

**DOI:** 10.1101/2021.05.05.442860

**Authors:** Guodong Wang, Qi Li, Junjie Xu, Shuai Zhao, Rui Zhou, Zhenkang Chen, Wentong Jiang, Xue Gao, Shuang Zhou, Zhiyu Chen, Quanzhi Sun, Chengyuan Ma, Lin Chen, Bihan Shi, Ying Guo, Haiyan Wang, Xia Wang, Huaiye Li, Tao Cai, Yibing Wang, Zhineng Chen, Fengchao Wang, Qinghua Liu

## Abstract

Classical forward and reverse mouse genetics approaches require germline mutations and, thus, are unwieldy to study sleep functions of essential genes or redundant pathways. It is also time-consuming to conduct electroencephalogram/electromyogram-based mouse sleep screening owning to labor-intensive surgeries and genetic crosses. Here, we describe a highly accurate SleepV (video) system and adeno-associated virus (AAV)-based adult brain chimeric (ABC)- expression/knockout (KO) platform for somatic genetics analysis of sleep in adult mice. A pilot ABC-expression screen identifies CREB and CRTC1, of which constitutive or inducible expression significantly reduces quantity and quality of non-rapid eye movement sleep. Whereas ABC-KO of exon 13 of *Sik3* by AAV-Cre injection in *Sik3-E13^flox/flox^* adult mice phenocopies *Sleepy (Sik3^Slp/+^)* mice, ABC-CRISPR of *Slp/Sik3* reverses hypersomnia of *Sleepy* mice, indicating a direct role of SLP/SIK3 kinase in sleep regulation. Multiplex ABC-CRISPR of both orexin/hypocretin receptors causes narcolepsy-like episodes, enabling one-step analysis of redundant genes in adult mice. Finally, ABC-expression/KO screen identifies Ankrd63 and NR1 as two potentially new sleep regulators. Therefore, this somatic genetics approach should facilitate high-throughput analysis of sleep regulatory genes, especially for essential or redundant genes, in adult mice by skipping the mouse development and genetic crosses.

## INTRODUCTION

Despite of significant advance in understanding the neural pathways that control executive sleep/wake switching (Liu and Dan, 2019; Saper et al., 2010; Saper et al., 2005; Scammell et al., 2017; Weber and Dan, 2016), the molecular mechanisms of mammalian sleep regulation are largely unclear (Allada et al., 2017; Sehgal and Mignot, 2011; Webb and Fu, 2021). It remains to be identified which genes constitute the core sleep regulatory pathways, and where these molecular pathways may function in the mouse brain. However, it is very costly and time-consuming to conduct large-scale mouse sleep screening for two main reasons: 1) both forward and reverse genetics approaches require germline mutations and genetic crosses (3 months/cross); 2) the electroencephalogram (EEG)/electromyogram (EMG)-based sleep analysis requires labor-intensive and invasive surgeries.

Accumulating evidence suggest that sleep is essential for survival in invertebrate and vertebrate animals (Bentivoglio and Grassi-Zucconi, 1997; Rechtschaffen et al., 1989; Shaw et al., 2002; Vaccaro et al., 2020). Thus, it is plausible that core sleep regulatory genes may be essential for survival in mice. It is estimated that about one third of ∼23,000 mammalian genes are essential genes (Dickinson et al., 2016), which often encode structural proteins, housekeeping enzymes, or signaling proteins with critical roles at multiple stages of development (Tian et al., 2018). However, classical mouse genetic approaches are unwieldy to study the sleep functions of essential genes owing to early lethality caused by germline mutations.

Conditional knockout (KO) mice are commonly used to bypass early lethality and analyze the temporal and/or tissue-specific functions of essential genes (Gierut et al., 2014). Typically, this strategy involves crossing of tissue-specific Cre recombinase or tamoxifen-dependent Cre^ERT2^ expressing transgenic mice with conditional flox mice that contain two loxP sites flanking a critical exon(s) of target gene (Feil et al., 1996; Gierut et al., 2014). The Cre/loxP-mediated site-specific recombination will exercise the critical exon(s) and disrupt the target gene in a tissue-specific and/or temporal manner. However, this time-consuming (1 to 2 years) approach requires construction of appropriate Cre transgenic and flox mouse strains, but also multiple genetic crosses to generate sufficient number of conditional KO mice for comprehensive sleep analysis (Gierut et al., 2014).

Given the vital importance of sleep in physiology and survival, it is likely that redundant pathways exist for sleep regulation. Thus, ablation of a gene of interest may cause mild or no sleep phenotype owning to genetic redundancy. Additionally, mice with germline mutations may adapt or compensate for sleep phenotypes before the EEG/EMG-based sleep analysis that is normally conducted in adult mice. Moreover, it is a costly and tedious process to generate double or triple KO mice by conventional germline genetics methods. A combination of triple-target CRISPR/Cas9 and modified embryonic stem cell technologies facilitates biallelic KO of redundant genes for sleep phenotype analysis in a single generation, but it does not work for essential genes (Sunagawa et al., 2016).

Recombinant adeno-associated viruses (AAVs) have been widely used as vehicles for gene expression, knockdown/knockout and gene therapy in the central nervous system (CNS) (Borel et al., 2014; Suzuki et al., 2016; Yang et al., 2016). To bypass the blood-brain barrier (BBB) and spatially restrict gene expression, these applications often require local AAV injection into specific regions of the mouse brain (Castro et al., 2021; Hunker et al., 2020; Zell et al., 2020). Alternatively, intravenous administration of AAVs provides a non-invasive strategy for systemic gene delivery into the CNS (Choudhury et al., 2016; Deverman et al., 2016; Foust et al., 2009). In particular, Gradinaru and colleagues have used the <u>C</u>re-<u>re</u>combination-based <u>A</u>AV-targeted <u>e</u>volution (CREATE) to isolate AAV9 variants, such as AAV-PHP.B and AAV-PHP.eB, which can efficiently bypass BBB and transduce the majority of adult brain neurons and astrocytes in certain mouse strains (Chan et al., 2017; Deverman et al., 2016). Both AAV-PHP.B and AAV-PHP.eB have been successfully used to deliver systemic gene expression (Liu et al., 2021; Luoni et al., 2020; Silva-Pinheiro et al., 2020), or target gene ablation by CRISPR/Cas9 across the adult mouse brain neurons (Torregrosa et al., 2021; Xiao et al., 2021; Yamaguchi and de Lecea, 2019).

Here, we systematically assembled, optimized and utilized this AAV-PHP.eB-based <u>a</u>dult <u>b</u>rain <u>c</u>himeric (ABC)-expression/KO platform for somatic genetics analysis of sleep in adult mice **(Figure 1)**. We also developed a highly accurate, non-invasive and fully automated SleepV (video) system for high-throughput mouse sleep screening. By skipping the mouse development and genetic crosses, this somatic genetics approach, coupled with the SleepV system, should facilitate highly efficient analysis of sleep regulatory genes, especially for essential or redundant genes, in adult mice.

**Figure 1.**
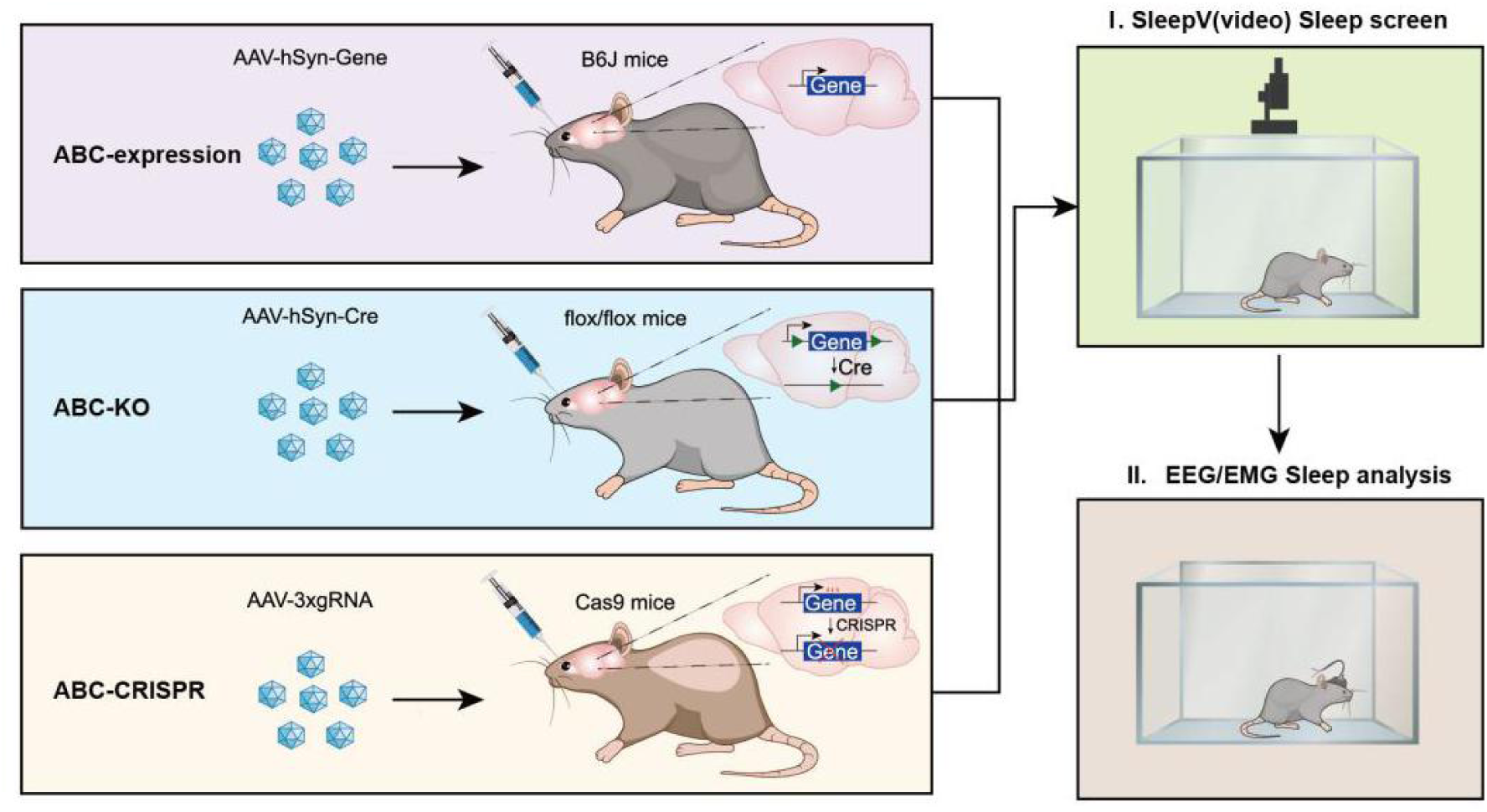
Adult brain chimeric (ABC) platform for somatic genetics analysis of sleep genes in adult mice. ABC-expression in C57BL/6J adult mice by retro-orbital injection of AAV-PHP.eB to deliver systemic gene expression from the human synapsin (hSyn) promoter in the adult brain neurons. ABC-KO of target gene in conditional flox/flox adult mice by retroorbital injection of AAV-PHP.eB expressing Cre recombinase. Multiplex ABC-CRISPR of target genes by retroorbital injection of AAV-3xsgRNA expressing a set of triple sgRNAs targeting different exons of the same gene. The ABC-expression/KO/CRISPR mice are subjected to high-throughput sleep analysis by SleepV (video) system followed (or directly) by EEG/EMG recording.

## RESULTS

### Development of ABC platform for somatic genetics analysis of sleep in adult mice

Sleep and wakefulness are two alternate physiological states of the brain, which globally impact the molecular, synaptic and cellular activities across the whole brain (Cirelli and Tononi, 1998; de Vivo et al., 2017; Diering et al., 2017; Elliott et al., 2014; Tononi and Cirelli, 2014; Wang et al., 2018). Thus, homeostatic sleep regulation likely involves the majority of neurons and possibly astrocytes across the adult mouse brain (Tononi and Cirelli, 2014; Wang et al., 2018). It has been reported that retro-orbital injection of 10^11^ vector genomes (vg) of single-stranded AAV-PHP.eB systemically transduced approximately 40% to 80% of adult brain neurons and astrocytes (Chan et al., 2017). Therefore, we hypothesized that AAV-PHP.eB-mediated ABC-expression or knockout of sleep regulatory genes should theoretically result in significant sleep phenotypes **(Figure 1)**.

To verify the efficiency of this AAV delivery system, we performed retro-orbital injection of 12-week old C57BL/6J mice with dual AAV-PHP.eB (10^12^ vg/mice), AAV-CBh-Cre and AAV-EF1α-DIO-H2B-eGFP **(Figure 1-figure supplement 1A)**. Thus, only brain cells co-transduced with both AAVs could exhibit the Cre-dependent expression of histone H2B-GFP fusion proteins. As shown by co-immunostaining of GFP and NeuN, a pan-neuronal marker protein **(Figure 1-figure supplement 1B and C)**, these two viruses efficiently co-transduced the majority of the adult brain neurons two weeks after AAV administration. Quantification of double positive cells showed high percentage of GFP-expressing neurons across nine different brain regions, ranging from averaging 56.6% in the hippocampus to 88.7% in the thalamus **(Figure 1-figure supplement 1D)**. More importantly, ABC-expression of the GFP reporter did not affect the sleep-wake architecture in the AAV-PHP.eB-injected mice relative to no virus injected control mice **(Figure 1-figure supplement 1E-H)**, suggesting that it is feasible to use the ABC-expression/KO platform for rapid and efficient somatic genetics analysis of sleep regulatory genes.

**Figure 1-figure supplement 1.**
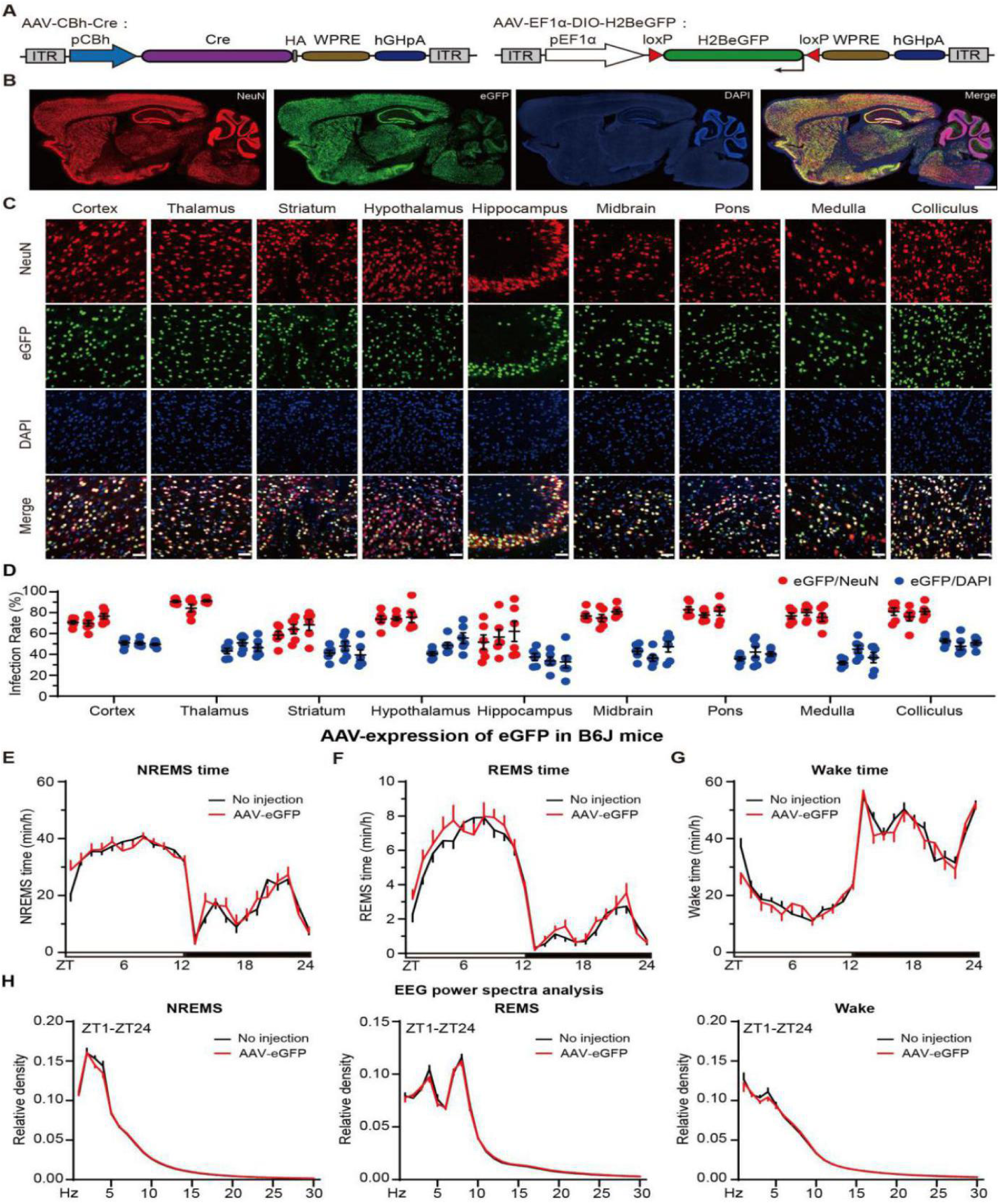
Development of ABC platform for somatic genetics analysis of sleep genes. **(A)** Schematic of the AAV-CBh:Cre and AAV-pEF1α:DIO-H2B-eGFP constructs. **(B)** Representative images showing co-immunostaining of eGFP and NeuN in the sagittal brain sections of AAV-CBh-Cre and AAV-EF1α-DIO-eGFP co-injected mice. **(C)** Representative images of NeuN^+^ (red) neurons or DAPI^+^ (blue) cells that also express eGFP in nine different brain regions. **(D)** Quantification of the viral transduction rates, which is calculated by the percentage of NeuN^+^ neurons (red) and DAPI^+^ (blue) cells that express eGFP, in nine brain regions shown in (C). **(E-H)** Hourly plots of NREMS (E), REMS (F) or Wake (G) time and EEG power spectra analysis of NREMS, REMS and Wake states (H) of no virus (n=19) or AAV-hSyn-eGFP (n=9) injected mice. Data are mean ± s.e.m. (E-H) Two-way ANOVA with Dunn’s multiple comparisons test. n.s. not significant.

### A highly accurate SleepV (video) system expedites mouse sleep screening

The EEG/EMG recording is the “Gold Standard” method for sleep/wake analysis in mammals (Lo et al., 2004; Weiergraber et al., 2005). Based on the different patterns of electrical signals, each short (4 to 20-s) epoch of EEG/EMG data is classified into one of three states: wakefulness (Wake), rapid eye movement sleep (REMS) or non-rapid eye movement sleep (NREMS). NREMS, which occupies ∼90% of total sleep time, is characterized by high percentage of the delta (1-4 Hz) power of EEG spectrum. The delta power of NREMS measures sleep quality or depth, which is often regarded as a good index of homeostatic sleep need (Franken et al., 2001).

However, this EEG/EMG method is not optimal for high-throughput mouse sleep screening because it requires labor-intensive and invasive surgery to implant electrodes into the skull and muscle and extensive recovery time from surgery before EEG/EMG recording. Moreover, the semi-automated sleep staging software only has ∼90% accuracy and requires intensive efforts to manually correct the annotation of several days of EEG/EMG data per mouse. On the other hand, a number of non-invasive sleep monitoring systems have been developed and utilized for mouse sleep analysis, including the infrared video recording system (Banks et al., 2020; Fisher et al., 2012; Pack et al., 2007), the piezoelectric system tracking mouse movement (Flores et al., 2007; Yaghouby et al., 2016) and the plethysmography system monitoring the respiration of mouse (Sunagawa et al., 2016).

Here, we developed an artificial intelligence (AI)-augmented video-based sleep monitoring system that we named SleepV, which used a novel pattern recognition algorithm for sleep/wake staging based on inactivity/activity of the mouse **(Figures 2A and Figure 2-figure supplement 1A-B)**. First, SleepV algorithm uses Gaussian filtering, adaptive and global thresholding to extract from every image frame (25 frames/sec) the suspected regions of interest (ROIs), which are then judged to be a mouse or not by a pre-trained deep neural network. Secondly, to determine whether the mouse is active or not, the algorithm calculates a high confidence difference score for the mouse between “t” and “t+m” frames by integrating the network prediction score (predict), the mouse mask area (Intersection of Union, IoU), the center of mass of the mask (activity), and the gray information (color) within the detected mask. Finally, SleepV defines the sleep state as ≥ 40- s of continuous immobility similar to previously reported video-based sleep monitoring systems (Banks et al., 2020; Fisher et al., 2012; Pack et al., 2007).

**Figure 2.**
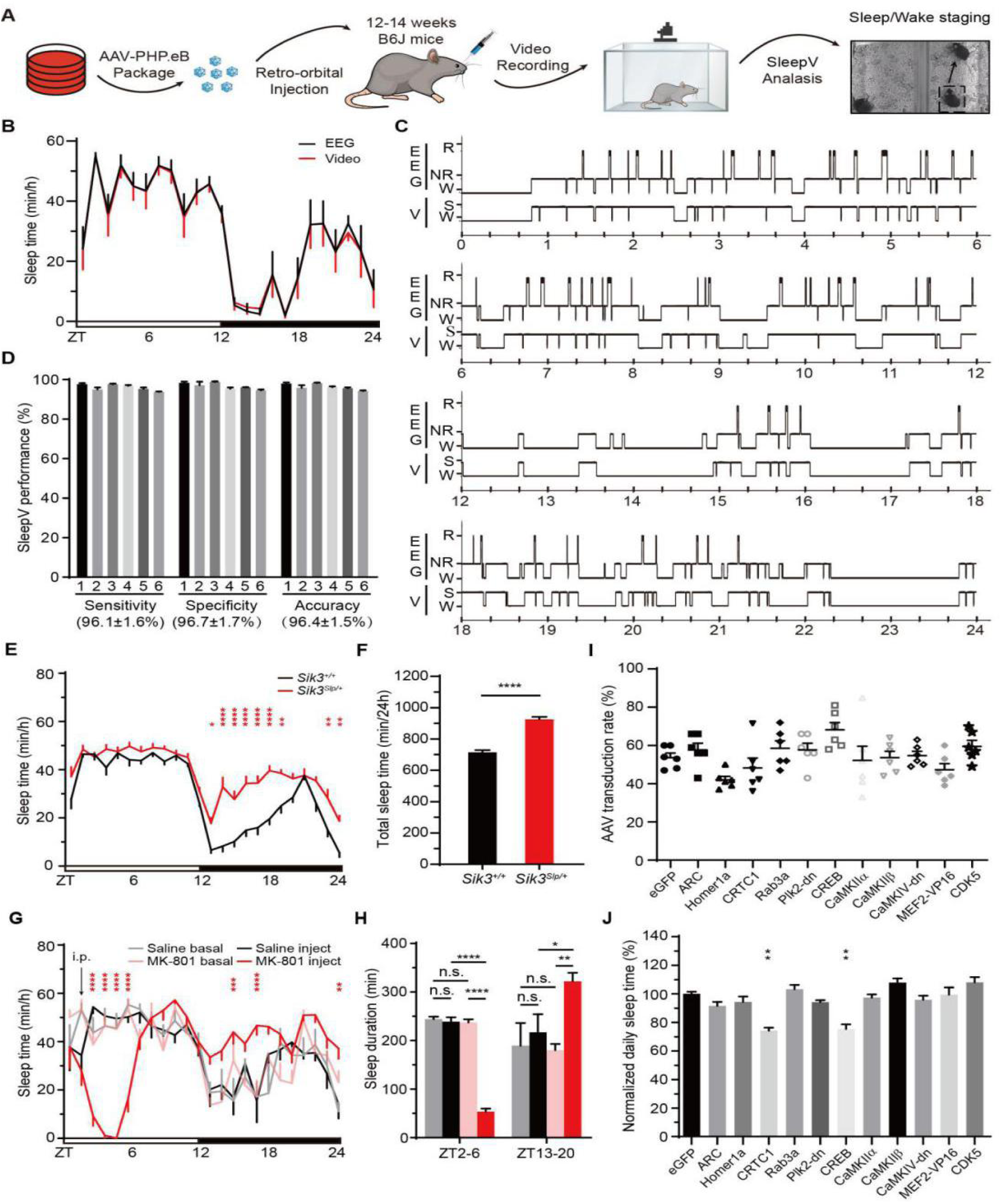
Development of a SleepV(video)-based high-throughput ABC sleep screening platform. **(A)** Schematic of video-based ABC-expression sleep screening platform. **(B)** Hourly plot of sleep time of six C57BL/6J mice by simultaneous SleepV(video) and EEG/EMG analysis. **(C)** Epoch-by-epoch comparison of sleep/wake staging of the same mouse by simultaneous SleepV and EEG/EMG analysis. **(D)** Quantitative analysis of the sensitivity, specificity and accuracy of sleep/wake staging of six mice by SleepV (total 77760 epochs, 20s/epoch). **(E)** Hourly plot of sleep time of *Sik3^+/+^* (n=11) and *Sik3^Slp/+^* (n=11) mice analyzed by SleepV. **(F)** Quantification of daily sleep time in *Sik3^+/+^* (n=11) and *Sik3^Slp/+^* (n=11) mice. **(G)** Hourly plot of sleep time of baseline condition or after i.p. injection of saline (black, n=4) or 2mg/kg MK-801 (red, n=4) at ZT2 in C57BL/6J mice. **(H)** Quantification of sleep time during ZT2-6 and ZT13-20 after saline or MK-801 injection. **(I)** Graph showing viral transduction rates of cortical neurons in AAV-hSyn-GeneX injected mice. **(J)** Graph showing daily sleep time of ABC-GeneX mice (n ≥ 5) normalized to ABC-eGFP mice. Data are mean ± s.e.m. (B, E, G, H) Two-way ANOVA with Dunn’s multiple comparisons test. (F) Unpaired t test. (J) One-way ANOVA with Dunn’s multiple comparisons test. n.s. not significant; * *p* < 0.05; ** *p* < 0.01; **** *p* < 0.0001.

**Figure 2-figure supplement 1.**
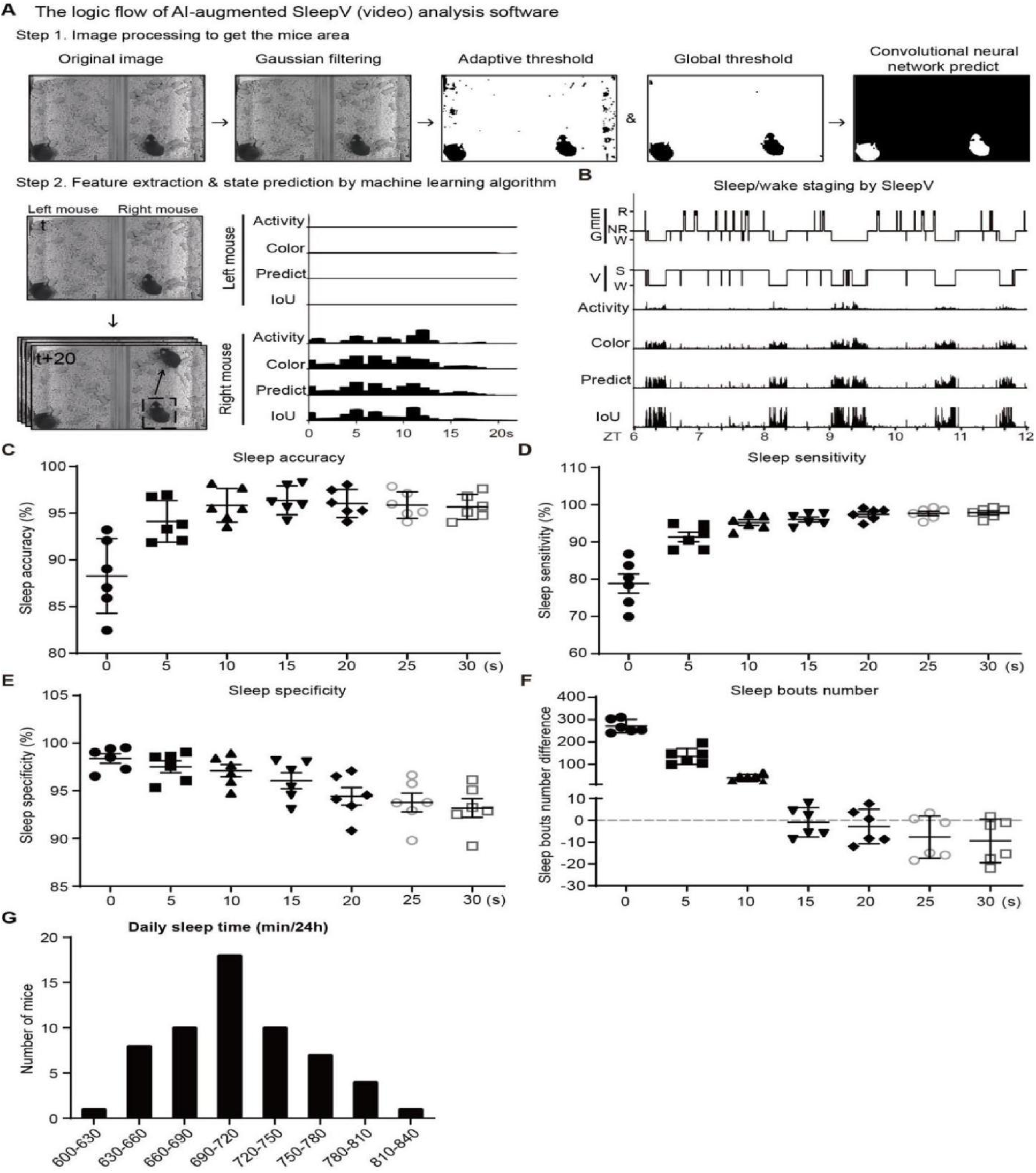
Development of a highly accurate SleepV system for high-throughout mouse sleep screening. **(A)** The logic flow of sleep/wake staging of video recording by SleepV software. **(B)** Epoch-by-epoch comparison of sleep/wake staging of the same mouse by simultaneous SleepV and EEG/EMG recording and analysis. **(C-F)** Graphs showing the accuracy, sensitivity, specificity and sleep bout number of sleep-wake staging by SleepV after filtering 0, 5, 10, 15, 20, 25 or 30s of small movements during sleep. **(G)** Distribution of daily sleep time in fifty-nine C57BL/6J mice analyzed by SleepV.

**Figure 2-Video S1.**

**SleepV can easily detect sleep/wake state transition.**

Shown are EEG/EMG characterized behavior states, SleepV characterized behavior states, and real time color&activity changes. NREM (N)-Wake (W)-NREM transitions were observed on the left mouse. REM (R)-Wake (W) transition was observed on the right mouse. The video is shown at 3x the original speed. S, sleep.

**Figure 2-Video S2.**

**SleepV can filter small movement during sleep.**

Shown are EEG/EMG characterized behavior state, SleepV characterized behavior state, and real time color&activity changes. Both the left and right mice showed small movement during NREM sleep, which could be filtered by SleepV. The video is shown at 3x the original speed.

We performed silmultaneous infrared video recording and EEG/EMG recording on six C57BL/6J adult mice to examine the accuracy of sleep/wake staging by fully automated SleepV analysis. **(Video S1)**. As compared to semi-automated EEG/EMG analysis with manual corrections, SleepV showed suboptimal accuracy (average 88.2%) by overestimating the number of sleep bouts owning to misjudging small mouse movements during sleep as waking **(Figure 2-figure supplement 1C and F)**. To improve the performance of SleepV, we systematically tested a series of thresholds of 0, 5, 10, 15, 20, 25, 30-s to filter subtle mouse movements during sleep, and compared the number of sleep bouts as well as the accuracy, sensitivity and specificity of sleep/wake staging **(Figure 2-figure supplement 1C-F and Video S2)**. Notably, we found that the best performance of SleepV was achieved with a 15-s threshold, that is, by annotating ≤ 15-s of activity between two sleep bouts as sleep.

With this improvement, the accuracy of sleep/wake staging by SleepV was now comparable to that of semi-automatic EEG/EMG analysis with human corrections **(Figure 2B)**. Although SleepV could not distinguish between NREMS and REMS, there was a remarkable 95-99% epoch-by-epoch agreement between the two methods, with ∼96% in sensitivity, specificity and accuracy of sleep/wake staging by SleepV **(Figure 2C and 2D)**. The mean accuracy (96.4%) of SleepV is significantly higher than that (92-94%) of previous reported video-based sleep monitoring systems (Banks et al., 2020; Fisher et al., 2012; Pack et al., 2007).

Next, we used SleepV to record the sleep/wake cycles of fifty-nine C57BL/6J adult mice for three consecutive days. The distribution of daily sleep time measured by SleepV in these mice was comparable to that of EEG/EMG analysis in previous studies **(Figure 2-figure supplement 1G**) (Funato et al., 2016; Wang et al., 2018). Moreover, SleepV could easily distinguish the hypersomnia phenotype of *Sleepy* (*Sik3^Slp/+^*) mice, which showed on average ∼210 min increase in daily sleep time relative to wild-type littermates **(Figure 2E and 2F)**. To test whether SleepV could detect dynamic changes in sleep/wake behavior, we intraperitoneally (i.p.) injected C57BL/6J mice with 2mg/kg MK-801–a specific inhibitor of NMDA receptor–at zeitgeber time 2 (ZT2) (Sunagawa et al., 2016; Tatsuki et al., 2016; Wang et al., 2018). As compared to the baseline and saline injection controls, MK-801 treatment rapidly decreased sleep time during ZT2-6, which was followed by rebound sleep during ZT13-20 **(Figure 2G and 2H)**. Collectively, these results demonstrate that SleepV is a highly accurate, non-invasive and fully automatic sleep monitoring system that is suitable for high-throughput mouse sleep screening.

### A pilot ABC-expression sleep screen of synaptic plasticity regulators

Accumulating studies suggest a close link between synaptic plasticity and sleep need regulation (Donlea et al., 2009; Ganguly-Fitzgerald et al., 2006; Huber et al., 2006; Huber et al., 2004). To test our hypothesis that changing synaptic plasticity could lead to changes in sleep need, we used SleepV system to conduct a pilot ABC-expression sleep screen of eleven known regulators of synaptic plasticity **(Figure 2I)**. These included the immediate early gene products ARC and Homer1a (Diering et al., 2017; Hu et al., 2010; Plath et al., 2006; Rial Verde et al., 2006; Shepherd et al., 2006), <u>c</u>yclin-<u>d</u>ependent <u>k</u>inase 5 (CDK5) (Bibb, 2003), Rab3a (Kapfhamer et al., 2002), dominant negative form of <u>p</u>olo-like <u>k</u>inase 2 (Plk2-dn) (Seeburg et al., 2008), <u>Ca</u>^2+^/cal<u>m</u>odulin-dependent protein <u>k</u>inase II (CaMKIIα/β) and IV (CaMKIV) (Ibata et al., 2008; Lisman et al., 2012), activity-dependent transcriptional factors <u>c</u>yclic AMP-<u>r</u>esponse <u>e</u>lement <u>b</u>inding protein (CREB) and <u>CREB r</u>egulated transcriptional <u>c</u>oactivator 1 (CRTC1), and constitutively active MEF2^VP16^–a fusion protein between the DNA binding domain of MEF2 and VP16 transactivation domain (Benito and Barco, 2010; Ch’ng et al., 2012; Flavell et al., 2006; Kandel, 2012; Nonaka et al., 2014). As shown by co-immunostaining of HA-tag and NeuN, intravenous administration of AAV-PHP.eB consistently resulted in systemic expression of target genes in 40-80% of neurons across the adult mouse brains **(Figure 2I)**. While ABC-expression of most genes had little effect on the sleep/wake cycle, this pilot screen identified two potential hits, CREB and CRTC1, which significantly reduced daily sleep amount **(Figure 2J)**.

CREB is a well-known transcriptional activator that binds as a dimer to the cAMP response elements (CRE), which contain a palindromic (TGACGTCA) or half-site (TGACG or CGTCA) sequence, in the promoter or enhancer regions of target genes (Comb et al., 1986; Montminy et al., 1986; Short et al., 1986). The *Creb1* gene encodes multiple CREB isoforms by alternative splicing, of which the α and Δ isoforms, but not the β isoform, show high affinity for CRE sites (Ruppert et al., 1992). Phosphorylation of CREB_α_ at serine 133 (or CREB_Δ_ at serine 119) by cAMP-dependent protein kinase (PKA) promotes the recruitment of coactivators, including histone acetyl transferases CBP/p300, and transcriptional activation of certain target genes (Chrivia et al., 1993; Gonzalez and Montminy, 1989; Shaywitz and Greenberg, 1999). Alternatively, CREB functions in tandem with CRTCs, also known as transducers <u>o</u>f <u>r</u>egulated <u>C</u>REB activity coactivators (TORCs), to activate the transcription of a specific subset of target genes (Conkright et al., 2003; Iourgenko et al., 2003).

### ABC-expression of CREB and/or CRTC1 reduces NREMS amount and delta power

We performed EEG/EMG recording to further characterize the sleep phenotypes caused by ABC-expression of CREB_Δ_. It has been shown that serine 133 to alanine (S133A) phosphor-mutation of CREB_α_ prevents transcriptional activation of specific target genes (Chrivia et al., 1993; Gonzalez and Montminy, 1989). Thus, we compared the sleep phenotypes as a result of ABC- expression of wild-type or S119A phosphor-mutant CREB_Δ_ in C57BL/6J adult mice. Co-immunostaining revealed that intravenous administration of AAV-hSyn-CREB_Δ_ or AAV-hSyn-CREB_Δ_ resulted in efficient transduction of the majority of cortical and thalamic neurons **(Figure 3A and 3B)**. Both ABC-CREB_Δ_ and ABC-CREB_Δ_^S119A^ mice, relative to ABC-eGFP mice, exhibited on average ∼120 min decrease in daily NREMS amount accompanied by reduced NREMS delta power, which occurred mostly during the dark phase **(Figure 3C-E and Figure 3-figure supplement 1A-E)**. These results suggest that ABC-expression of CREB_Δ_ reduces NREMS amount and NREMS delta power in a manner independent of S119 phosphorylation.

**Figure 3.**
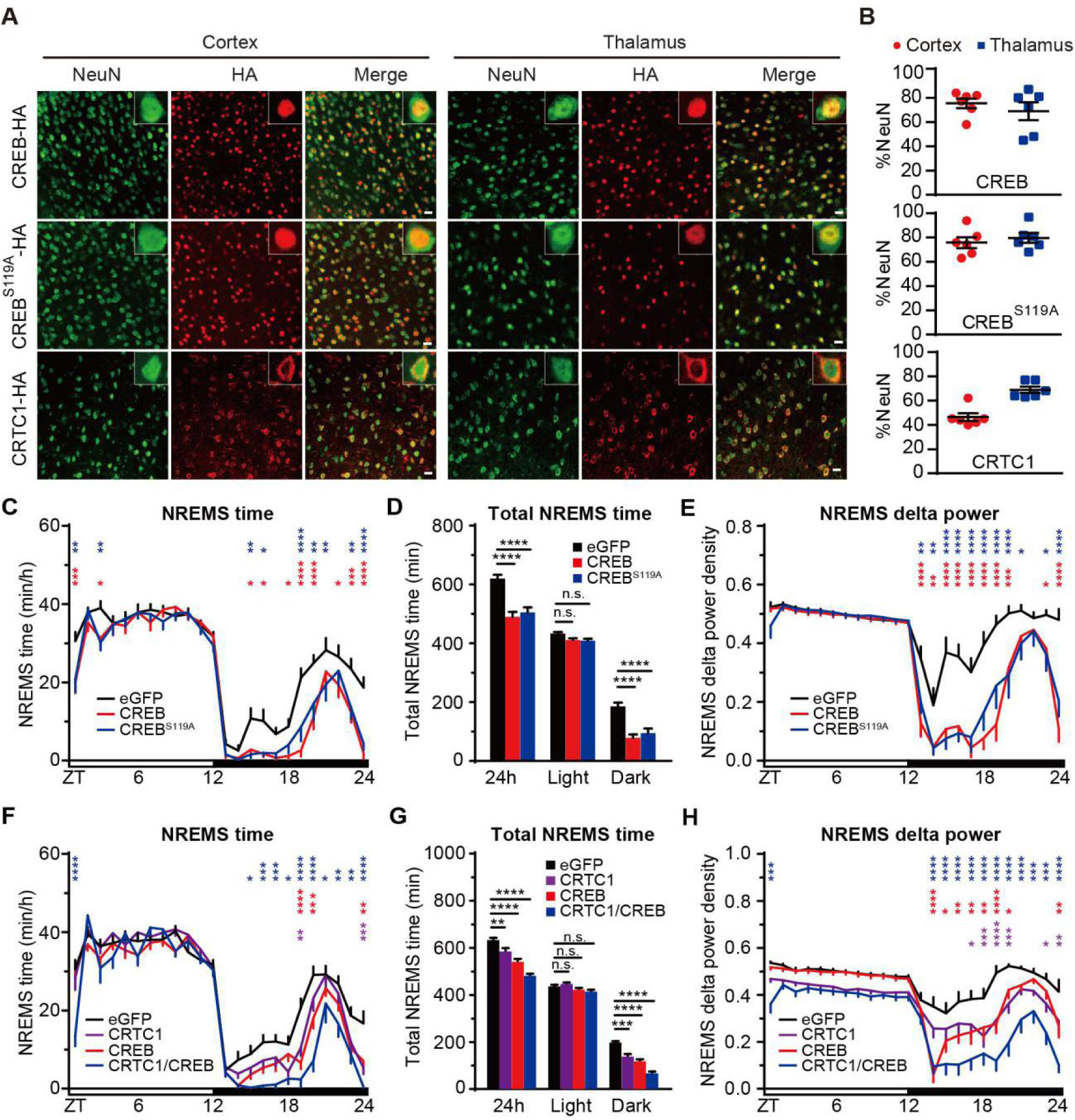
ABC-expression of CREB and/or CRTC1 reduces NREMS amount and delta power. **(A)** Co-immunostaining of HA^+^ (red) and NeuN^+^ (green) neurons in the cortex and thalamus of AAV-hSyn-CREB (ABC-CREB), AAV-hSyn-CREB^S119A^ (ABC-CREB^S119A^) and AAV-hSyn-CRTC1 (ABC-CRTC1)-injected mice. **(B)** Quantification of the viral transduction rates, which is calculated by the percentage of NeuN^+^ neurons that express HA- tagged proteins, in the cortical and thalamic neurons of ABC-CREB, ABC-CREB^S119A^ and ABC-CRTC1 mice. **(C-E)** Hourly plot of NREMS time (C), quantification of total NREMS time (D) and hourly plot of NREMS delta power (E) in the ABC-eGFP (n=11), ABC-CREB (n=12) and ABC-CREB^S119A^ (n=11) mice. Shown above are the statistical analysis for comparison between ABC-CREB (red*) or ABC-CREB^S119A^ (blue*) mice and control ABC-eGFP mice. **(F-H)** Hourly plot of NREMS time (F), quantification of total NREMS time (G) and hourly plot of NREMS delta power (H) in the ABC-eGFP, (n=12), ABC-CRTC1 (n=15), ABC-CREB (n=15) and ABC-CRTC1/CREB (n=12) mice. Shown above are statistical analysis for comparison between ABC-CRTC1 (purple*), ABC-CREB (red*), or ABC-CRTC1/CREB (blue*) mice and control ABC-eGFP mice. Data are mean ± s.e.m. (C-H) Two-way ANOVA with Dunn’s multiple comparisons test. n.s. not significant; * *p* < 0.05; ** *p* < 0.01; *** *p* < 0.001; **** *p* < 0.0001.

Next, we asked whether ABC-coexpression of CREB_Δ_ and CRTC1 could result in additive sleep phenotypes as compared to ABC-expression of either CREB_Δ_ or CRTC1 alone. In contrast to nuclear localization of CREB_Δ_, CRTC1 was predominantly localized in the cytoplasm **(Figure 3A)**. Relative to ABC-eGFP mice, ABC-CRTC1 mice exhibited ∼47 min decrease in daily NREMS time and reduced NREMS delta power during the dark phase **(Figure 3F-H and Figure 3-figure supplement 1F-J)**. It should be noted that SleepV overestimated the reduction of sleep time in ABC-CRTC1 mice by misjudging excessive muscle twitching during sleep as waking in these mice **(Figure 3-figure supplement 1K-L and Video S3)**. Importantly, ABC-coexpression of CREB_Δ_ and CRTC1 resulted in enhanced sleep phenotypes: ∼150 min decrease in daily NREMS time accompanied by lower NREMS delta power during the dark phase **(Figure 3F-H)**. These results suggest that the function of CREB in sleep regulation is probably facilitated by a CRTC-dependent mechanism. Furthermore, this pilot screen demonstrates the proof-of-principle that the combination of ABC platform and SleepV system can facilitate high-throughput screening of sleep genes in mice.

**Figure 3-figure supplement 1.**
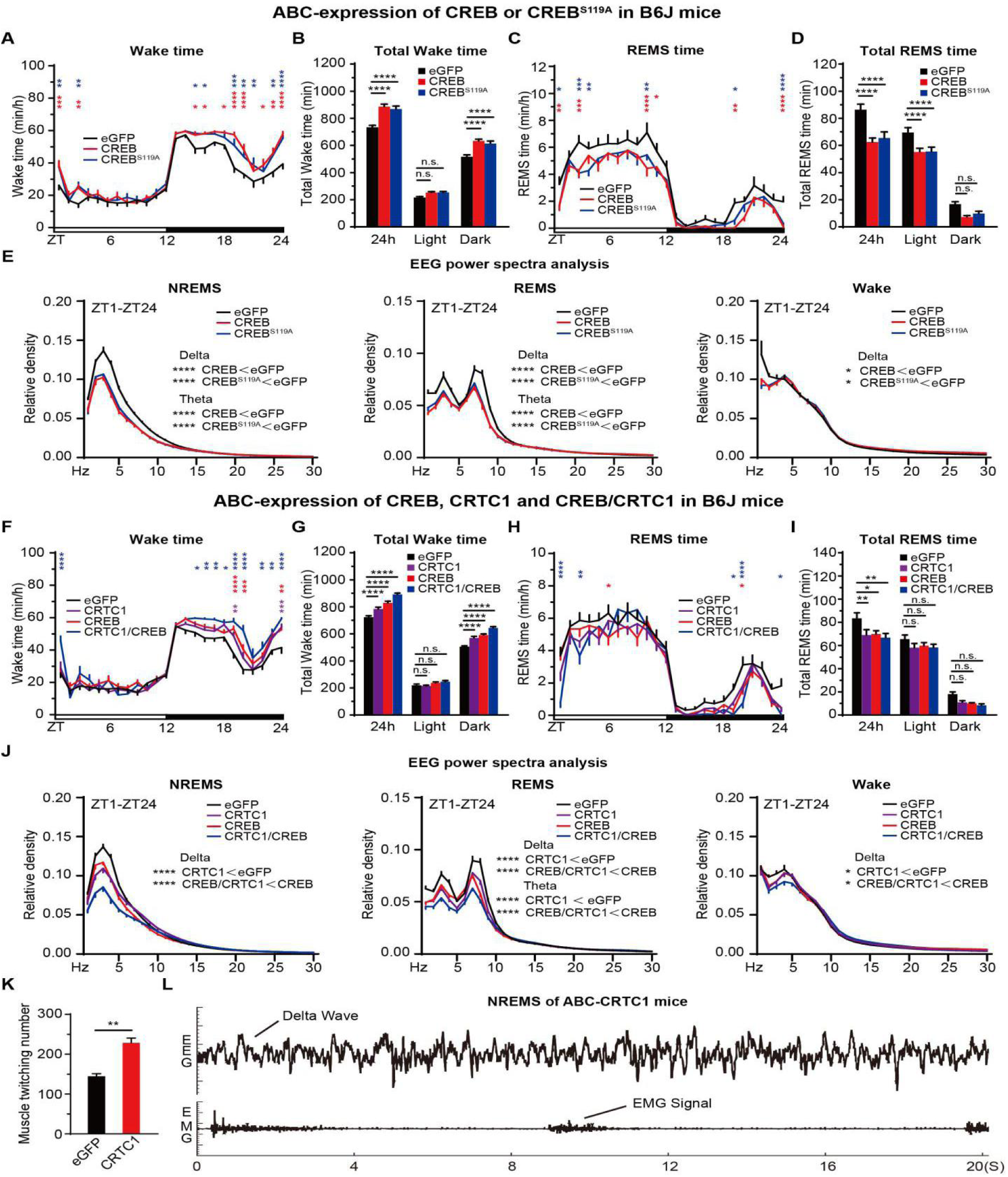
ABC-expression of CREB and/or CRTC1 reduces NREMS amount and delta power. **(A-E)** Hourly plots of Wake (A) or REMS (C) time, quantification of total Wake (B) or REMS (D) time and EEG power spectra analysis of NREMS, REMS and Wake states (E) in the ABC-eGFP (n=11), ABC-CREB_Δ_ (n=12) and ABC-CREB_Δ_ (n=11) mice. Shown above is statistical analysis for comparison between ABC-CREB (red*) or ABC-CREB^S119A^ (blue*) mice and control ABC-eGFP mice. **(F-J)** Hourly plots of Wake (F) or REMS (H) time, quantification of total Wake (G) or REMS (I) time and EEG power spectra analysis of NREMS, REMS and Wake states (J) in ABC-eGFP (n=12), ABC-CRTC1 (n=15), ABC-CREB_Δ_ (n=15) and ABC-CRTC1/CREB_Δ_ (n=12) mice. Shown above is the statistical analysis for comparison between ABC-CRTC1 (purple*), ABC-CREB_Δ_ (red*), or ABC-CRTC1/CREB_Δ_ (blue*) mice and control ABC-eGFP mice. **(K)** Quantification of muscle twitching episodes of ABC-eGFP and ABC-CRTC1 mice during NREMS. **(L)** Representative EEG/EMG hypnogram depicting frequent muscle twitching during NREMS in the ABC-CRTC1 mice. Data are mean ± s.e.m. (A-J) Two-way ANOVA with Dunn’s multiple comparisons test. (K) Unpaired t test. n.s. not significant;* *p* < 0.05; ** *p* < 0.01; *** *p* < 0.001; **** *p* < 0.0001.

**Figure 3-Video S3.**

**ABC-CRTC1 mouse shows frequent muscle twitching during sleep.**

### Inducible ABC-expression of CREB^VP16^ and/or CRTC1^CA^ causes strong sleep phenotypes

ABC-expression of constitutively active CREB^VP16^, a fusion protein between the DNA- binding domain of CREB and VP16 transactivation domain (Barco et al., 2002), or constitutively active CRTC1^CA^ containing two (S151A and S245A) phosphor-mutations (Sonntag et al., 2017), resulted in lethality one week after AAV injection in C57BL/6J adult mice. To study the immediate sleep phenotypes resulted from ABC-expression of CREB^VP16^ or CRTC1^CA^, we used a Tet-on inducible system to express CREB^VP16^ or CRTC1^CA^ in the adult mouse brains by co-injection of two AAV-PHP.eB viruses expressing rtTA from the EF1α promoter and CREB^VP16^ or CRTC1^CA^ from the TRE promoter, respectively **(Figure 4A)**. There was no difference in the baseline sleep-wake architecture among the inducible (i)ABC-eGFP, iABC-CREB^VP16^ and iABC-CRTC1^CA^ mice when transcription from the TRE promoter was inactive in the absence of Doxycycline (Dox) (data not shown). On the other hand, the expression of GFP, CREB^VP16^ or CRTC1^CA^ was rapidly induced in the brain cells of these mice within three days after drinking Dox-containing water **(Figure 4B, C and Figure 4-figure supplement 1A, B)**. Interestingly, a slight circadian shift of the sleep/wake cycle was observed at the light/dark transition among all mice possibly due to the effects of Dox **(Figure 4D-O)**. While ABC-induction of GFP did not affect total sleep/wake time, ABC-induction of CREB^VP16^ or CRTC1^CA^ caused progressive decrease in daily amounts of NREMS and REMS accompanied by corresponding increase in total wake time during three days of Dox treatment **(Figure 4D-O and Figure 4-figure supplement 1C-N)**. Remarkably, ABC-CRTC1^CA^ mice were almost constantly awake on day 3 of Dox treatment as shown by 91.8% and 97.1% reduction in NREMS and REMS, respectively **(Figure 4L-O and Figure 4-figure supplement 1K-N)**. These results suggest that iABC-expression system can be used to study the immediate sleep phenotypes of target genes of which constitutive expression causes rapid lethality in adult mice.

**Figure 4.**
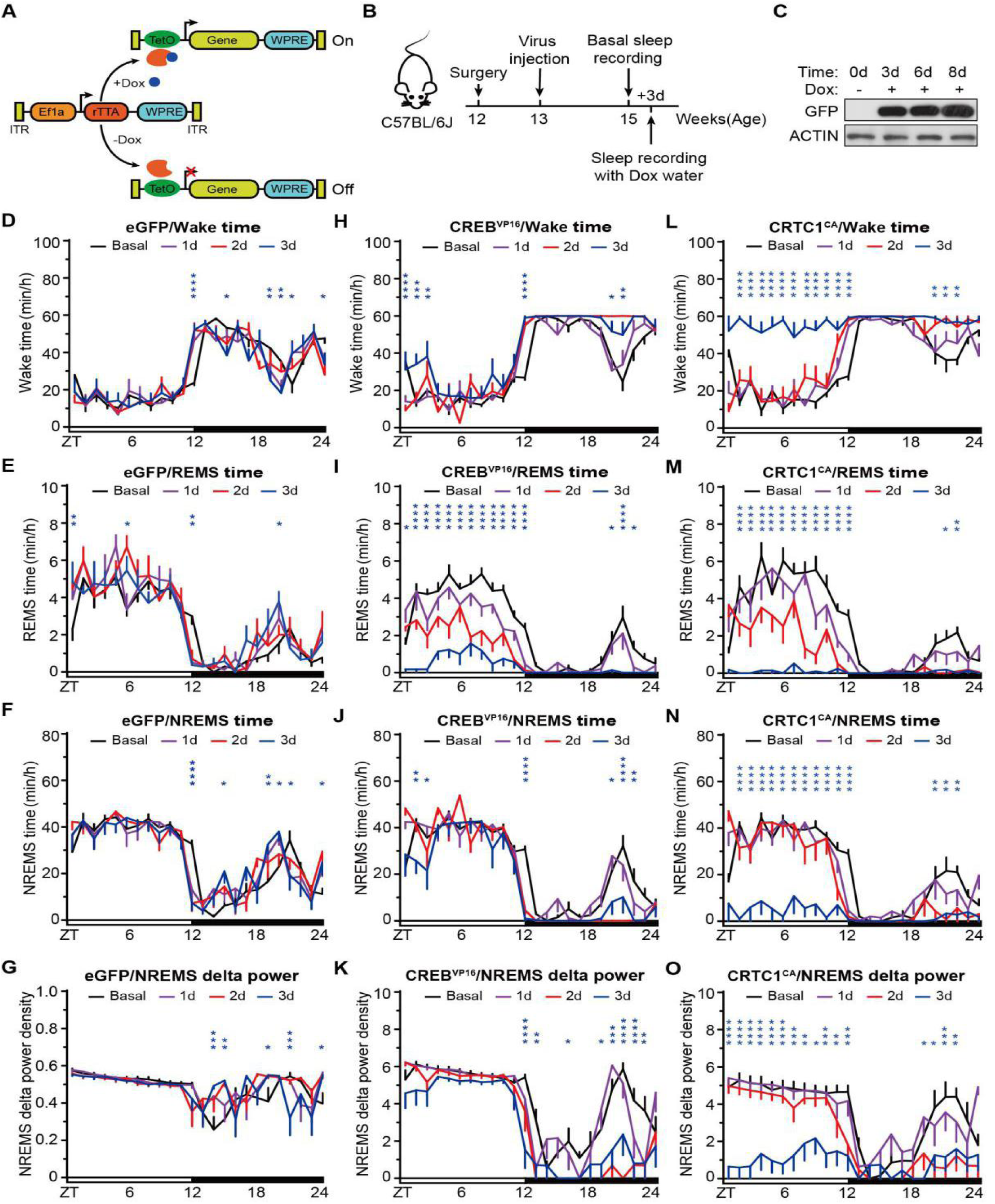
Inducible ABC-expression of CREB^VP16^ or CRTC1^CA^ causes significant sleep phenotypes. **(A)** Schematic of Tet-on inducible (i)ABC-expression system with or without Doxcycline (Dox). **(B)** A flow chart of the iABC-expression and EEG/EMG sleep recording experiment. **(C)** Immunoblotting of whole brain lysates from iABC-eGFP mice before and after Dox treatment with anti-GFP and anti-ACTIN antibodies. **(D-G)** Hourly plots of Wake time (D), REMS time (E), NREMS time (F) and NREMS delta power (G) of the iABC-eGFP mice (n=8). **(H-K)** Hourly plots of Wake time (H), REMS time (I), NREMS time (J) and NREMS delta power (K) in the iABC-CREB^VP16^ mice (n=7). **(L-O)** Hourly plots of Wake time (L), REMS time (M), NREMS time (N) and NREMS delta power (O) in the iABC-CRTC1^CA^ mice (n=7). Data are mean ± s.e.m. (D-O) Two-way ANOVA with Dunn’s multiple comparisons test. n.s. not significant; * *p* < 0.05; ** *p* < 0.01; *** *p* < 0.001; **** *p* < 0.0001.

**Figure 4-figure supplement 1.**
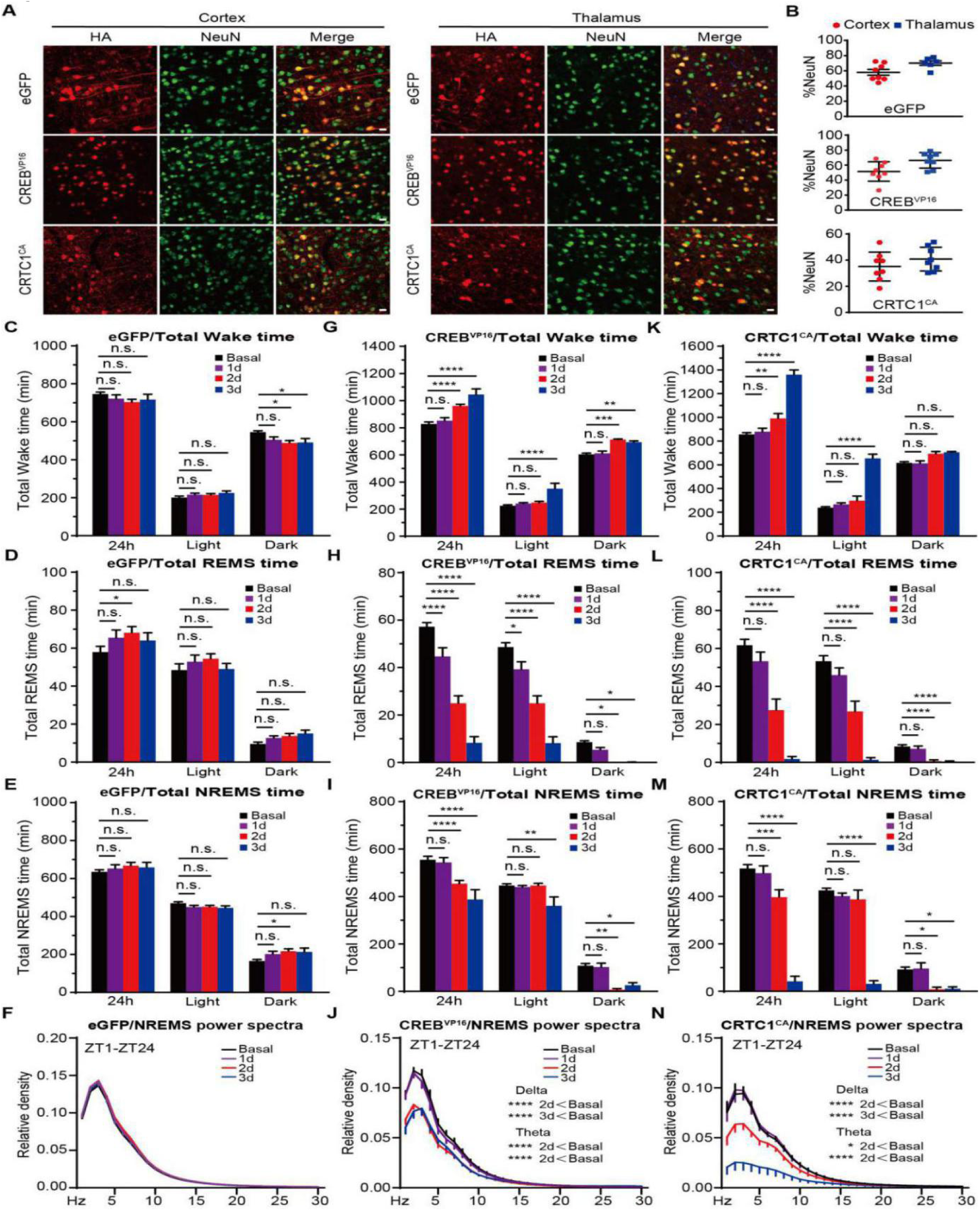
Inducible ABC-expression of CREB^VP16^ or CRTC1^CA^ causes significant sleep phenotypes. **(A)** Co-immunostaining of HA^+^ (red) and NeuN^+^ (green) neurons in the cortex and thalamus of the the inducible (i) ABC-eGFP, ABC-CREB^VP16^ and ABC-CRTC1^CA^ mice. Quantification of the viral transduction rates, which is calculated by the percentage of NeuN^+^ neurons that express HA-tagged proteins, in the cortical and thalamic neurons showed in (A). **(C-F)** Quantification of total Wake time (C), REMS time (D), or NREMS time (E) and EEG power spectra analysis of NREMS (F) in the iABC-eGFP mice (n=8). **(G-J)** Quantification of total Wake time (G), REMS time (H), or NREMS time (I) and EEG power spectra analysis of NREMS (J) in the iABC-CREB^VP16^ mice (n=7). **(K-N)** Quantification of total Wake time (K), REMS time (L), or NREMS time (M) and EEG power spectra analysis of NREMS (N) in the iABC-CRTC1^CA^ mice (n=7). Data are mean ± s.e.m. (C-N) Two-way ANOVA with Dunn’s multiple comparisons test. n.s. not significant; * *p* <0.05; ** *p* < 0.01; *** *p* < 0.001; **** *p* < 0.0001.

### ABC-KO of *Creb1* by Cre/loxP recombination increases daily NREMS amount

*Creb1* is an essential gene of which complete ablation results in perinatal lethality in mice (Bleckmann et al., 2002). A partial *Creb1* knockout strain, in which the α and Δ isoforms of CREB are deleted, is homozygous viable and exhibits ∼100 min increase of daily NREMS time (Graves et al., 2003). Moreover, forebrain-specific knockout of *Creb1* in the excitatory neurons similarly increases daily NREMS amount (Wimmer et al., 2021). To generate ABC-*Creb1^KO^* mice, we retro-orbitally injected *Creb1^flox/flox^* mice with AAV-PHP.eB expressing mCherry or Cre recombinase from the pan-neuronal hSyn promoter, respectively. Immunoblotting revealed that the level of CREB expression was reduced by ∼50% in whole brain lysates of AAV-hSyn-Cre injected mice relative to AAV-hSyn-mCherry injected mice **(Figure 5A-C)**. The efficiency of ABC- *Creb1^KO^* might be underestimated because CREB was also expressed in the astrocytes that could not be targeted by neuron-specific Cre expression (Pardo et al., 2017). Consistent with previous studies (Graves et al., 2003; Wimmer et al., 2021), ABC-*Creb1^KO^* mice exhibited ∼100 min increase in daily NREMS amount, with no significant change in NREMS delta power **(Figure 5D-F and Figure 5-figure supplement 1A-E)**. These results suggest that ABC-KO of *Creb1* can result in a significant sleep phenotype comparable to that of germline *Creb1* mutant mice (Graves et al., 2003; Wimmer et al., 2021).

**Figure 5.**
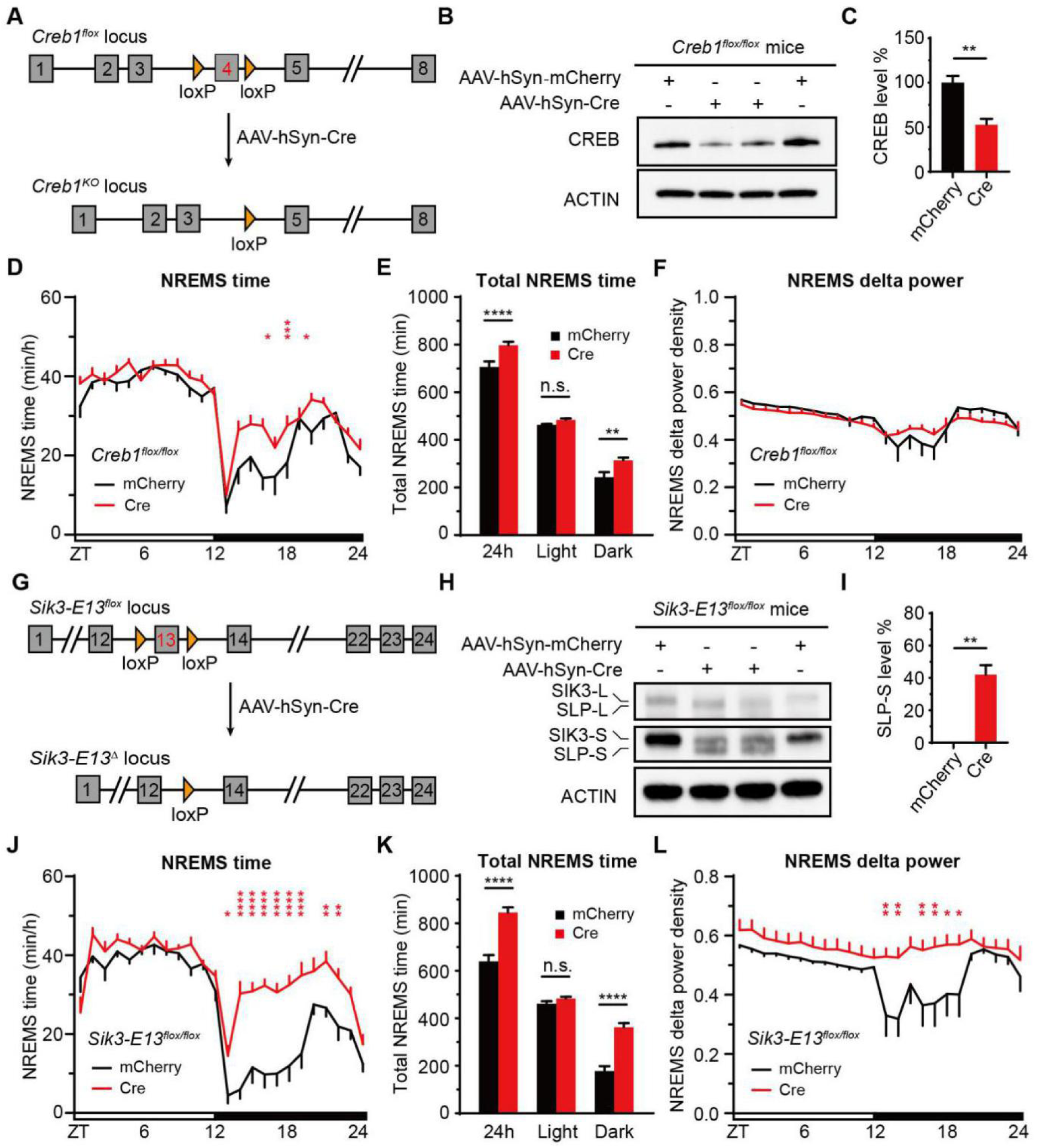
ABC-KO of *Creb1* or exon 13 of *Sik3* by Cre-loxP recombination causes hypersomnia. **(A)** Schematic of ABC-KO of *Creb1* by AAV-hSyn-Cre injection of *Creb1^flox/flox^* mice. **(B)** Immunoblotting of brain lysates from AAV-hSyn-mCherry or AAV-hSyn-Cre injected *Creb1^flox/flox^* mice with anti-CREB and anti-ACTIN antibodies. **(C)** Quantification of CREB expression in (B) (n=4). **(D-F)** Hourly plot of NREMS time (D), quantification of total NREMS time (E) and hourly plot of NREMS delta power (F) in the AAV-hSyn-mCherry (n=9) or AAV-hSyn-Cre (n=14) injected *Creb1^flox/flox^* mice. **(G)** Schematic of ABC-KO of exon 13 of *Sik3* by AAV-hSyn-Cre injection of *Sik3-E13^flox/flox^* mice. **(H)** Immunoblotting of brain lysates from AAV-hSyn-mCherry or AAV-hSyn-Cre injected *Sik3-E13^flox/flox^* mice with anti-SIK3 and anti-ACTIN antibodies. **(I)** Quantification of SLP-S expression in (H) (n=4), which is calculated by the percentage of SLP-S/(SIK3-S+SLP-S). **(J-L)** Hourly plot of NREMS time (J), quantification of total NREMS time (K) and hourly plot of NREMS delta power (L) in the AAV-hSyn-mCherry (n=8) or AAV-hSyn-Cre (n=8) injected *Sik3-E13^flox/flox^* mice. Data are mean ± s.e.m. (C and I) Unpaired t test. (D-F) and (J-L) Two-way ANOVA with Dunn’s multiple comparisons test. n.s. not significant; * *p* < 0.05; ** *p* < 0.01; *** *p* < 0.001; **** *p* < 0.0001.

**Figure 5-figure supplement 1.**
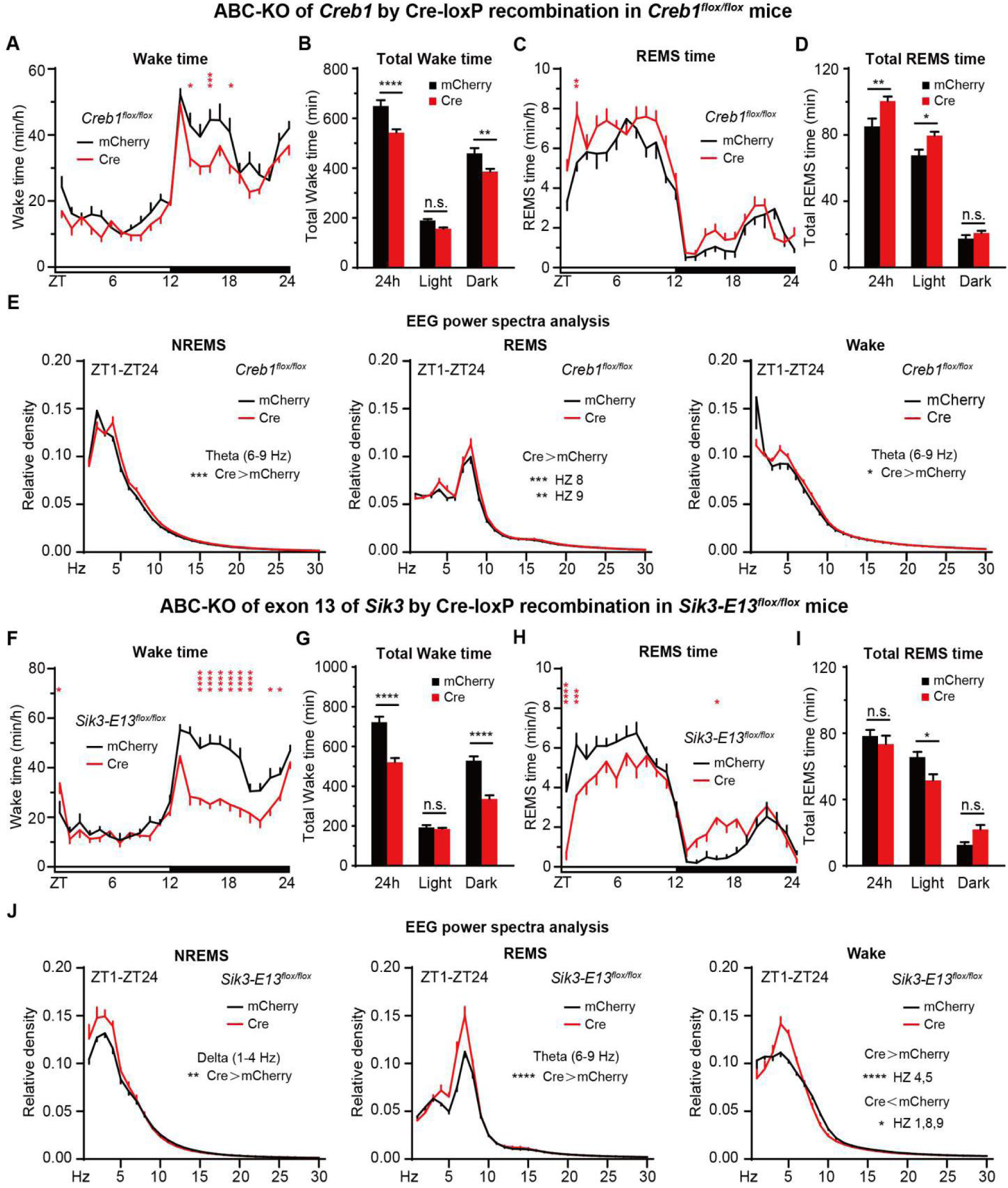
ABC-KO of *Creb1* or exon 13 of *Sik3* by AAV-mediated Cre-loxP recombination causes hypersomnia. **(A-E)** Hourly plots of Wake (A) or REMS (C) time, quantification of total Wake (B) or REMS (D) time and EEG power spectra analysis of NREMS, REMS and Wake states (E) in the AAV-hSyn-mCherry ( n=9) or AAV-hSyn-Cre ( n=14) injected *Creb1^flox/flox^* mice. **(F-J)** Hourly plots of Wake (F) or REMS (H) time, quantification of total Wake (G) or REMS (I) time and EEG power spectra analysis of NREMS, REMS and Wake states (J) in the AAV-hSyn-mCherry (n=8) or AAV-hSyn-Cre (n=8) injected *Sik3-E13^flox/flox^* adult mice. Data are mean ± s.e.m. (A-J) Two-way ANOVA with Dunn’s multiple comparisons test. n.s. not significant; * *p* <0.05; ** *p* < 0.01; *** *p* < 0.001; **** *p* < 0.0001.

### ABC-KO of exon 13 of *Sik3* phenocopies *Sleepy* mice

Forward genetic screening identified a *Sleepy* (*Sik3^Slp/+^*) mouse strain, in which a gain-of-function splicing mutation causes the skipping of exon 13 from *Sik3* transcripts and in-frame deletion of 52 amino acids from SIK3 proteins, an AMP-activated protein kinase (AMPK)-related protein kinase (Funato et al., 2016). The *Sik3^Slp/+^* mice exhibit ∼250 min increase in daily NREMS time with constitutively elevated NREMS delta power (Funato et al., 2016) as well as other phenotypes, such as obesity and reproductive defects (Funato and Yanagisawa, personal communications). Moreover, *Sik3* is an essential gene that is broadly expressed in mouse brain neurons (Funato et al., 2016). Thus, it remains unclear whether the hypersomnia of *Sik3^Slp/+^* mice is the primary phenotype owing to direct effects of SLP kinases in sleep regulation, or secondary phenotype resulted from the developmental defects of the brain or dysfunctions of peripheral organs.

To distinguish among these possibilities, we performed retro-orbital injection of AAV-hSyn-Cre into *Sik3-E13^flox/flox^* adult mice, such that Cre/loxP-mediated recombination would convert *Sik3-E13^flox^* into a functionally equivalent *Slp* (*Sik3-E13^Δ^*) allele **(Figure 5G)**. Immunoblotting estimated that mutant SLP proteins were expressed in at least 40% of the adult brain neurons following AAV-hSyn-Cre injection **(Figure 5H and 5I)**. Accordingly, ABC-KO of exon 13 of *Sik3* induced marked hypersomnia–200 to 300 min increase in daily NREMS time accompanied by constitutively elevated NREMS delta power–similar to that of *Sleepy* mice carrying the germline *Slp* mutation **(Figure 5J-L and Figure5-figure supplement 1F-J)**. These results suggest that the hypersomnia of *Sleepy* mice is the primary phenotype as a result of direct effects of mutant SLP kinases on the sleep regulatory machinery in the adult brain neurons.

### ABC-KO of genes by triple-target CRISPR in Cas9 mice

We reasoned that it would be more direct and faster to generate ABC-KO mice with the use of CRISPR/Cas9 technology. In this system, Cas9 nuclease is directed by single-guide (sg)RNA to introduce site-specific DNA break in the target gene, which is repaired by the error-prone non-homologous end-joining pathways, resulting in indel mutations (e.g., short deletions or insertions) (Hsu et al., 2014; Jinek et al., 2012; Wang et al., 2013). However, these Cas9-mediated indel mutations occur at a moderate frequency and not all mutations can ablate target gene function. Because the vast majority of adult brain neurons are non-dividing, terminally differentiated cells, the efficiency of ABC-KO by CRISPR needs to be nearly 100%, such that both alleles of the target gene are disrupted in almost all AAV-transduced brain cells. Although it was recently reported that intravenous injection of AAV-PHP.eB expressing single sgRNA can efficiently disrupt target gene in the majority of adult brain neurons (Xiao et al., 2021), this strategy often required extensive screening of sgRNAs and was not suitable for high-throughput analysis.

Multiplexing strategies using several sgRNAs targeting the same gene have been utilized to improve the efficiency of KO by CRISPR in various model organisms (Port et al., 2020; Xie et al., 2015; Yin et al., 2015). Notably, triple-target CRISPR in zygotes can produce whole-body biallelic knockout mice with 96-100% efficiency in a single generation (Sunagawa et al., 2016; Tatsuki et al., 2016). Thus, we compared the efficiency of ABC-KO by CRISPR by injecting *Rosa26^LSL-Cas9^* mice with AAV-PHP.eB expressing HA-tagged Cre recombinase from the hSyn promoter as well as one, two or three U6:sgRNA cistrons targeting *NeuN*, a ubiquitously expressed gene in the adult brain neurons **(Figure 6-figure supplement 1A and B)**. The efficiency of ABC-KO of *NeuN* was significantly higher with triple sgRNAs than with single or double sgRNAs **(Figure 6-figure supplement 1C)**. Moreover, the efficiency of ABC-*NeuN^KO^* increased in a viral dose-dependent manner and peaked at three weeks after AAV injection **(Figure 6-figure supplement 1D and E)**. Whole genome sequencing of ABC-*NeuN^KO^* mouse brain revealed on-target indel mutations and large inter-exon deletions, but rarely off-target mutations **(Figure 6-figure supplement 1F and G)**. Accordingly, immunoblotting showed that the level of NeuN expression was specifically reduced by ∼70%, whereas another pan-neuronal protein Tublin J remain unchanged in the ABC-*NeuN^KO^* brain lysates **(Figure 6-figure supplement 1H and I)**. Co-immunostaining of NeuN and Cre indicated that the expression of NeuN disappeared, indicative of biallelic KO of *NeuN*, in the majority of the adult brain neurons expressing both Cre and 3xsgRNA^NeuN^ **(Figure 6-figure supplement 1J)**.

### ABC-CRISPR of *Slp/Sik3* rescues hypersomnia of *Sik3^Slp/+^* mice

Because ABC-KO of exon 13 of *Sik3* could induce hypersomnia in adult mice **(Figure 5J-L)**, we hypothesized that ABC-KO of *Slp/Sik3* by triple-target CRISPR should rescue hypersomnia of *Sik3^Slp/+^* mice. To test our hypothesis, we performed ABC-CRISPR of *Slp/Sik3* alleles by injecting constitutively Cas9-expressing *Sik3^Slp/+^*;*Rosa26^Cas9/+^* adult mice with AAV-3xsgRNA*^Sik3^* expressing triple sgRNAs targeting different exons of *Sik3* gene **(Figure 6A)**. Whole genome sequencing and genomic PCR revealed both on-target indel mutations and large inter-exon deletions between distinct sgRNA target sites, but rarely off-target mutations, in the AAV-3xsgRNA^Sik3^ injected mouse brains **(Figure 6B and 6C)**. As shown by non-linear enhanced chemilumilescent (ECL) Western blotting, the levels of SIK3/SLP proteins were estimated to reduce by ∼75% in the ABC-*Sik3^KO^* brain lysates **(Figure 6D and 6E)**. By contrast, linear Western blotting, with the use of alkaline phosphatase (AP)-conjugated secondary antibodies, indicated that the levels of SIK3/SLP proteins were actually reduced by ∼50% in the ABC-*Sik3^KO^* brain lysates **(Figure6-figure supplement 2A-D)**. Importantly, ABC-CRISPR of *Slp/Sik3* resulted in ∼150 min reduction in daily NREMS time accompanied by constitutively reduced NREMS delta power in *Sik3^Slp/+^*;*Rosa26^Cas9/+^* male mice **(Figure 6F-H and Figure 6-figure supplement 2E-I)**. Likewise, ABC-CRISPR of *Slp/Sik3* using a second set of sgRNAs caused ∼180 min reduction in daily NREMS time with diminished NREMS delta power in *Sik3^Slp/+^*;*Rosa26^Cas9/+^* female mice **(Figure 6I-K and Figure 6-figure supplement 2J-N)**. These results suggest that the hypersomnia of *Sleepy* mice requires continuous expression of mutant SLP kinases in the adult brain neurons.

**Figure 6.**
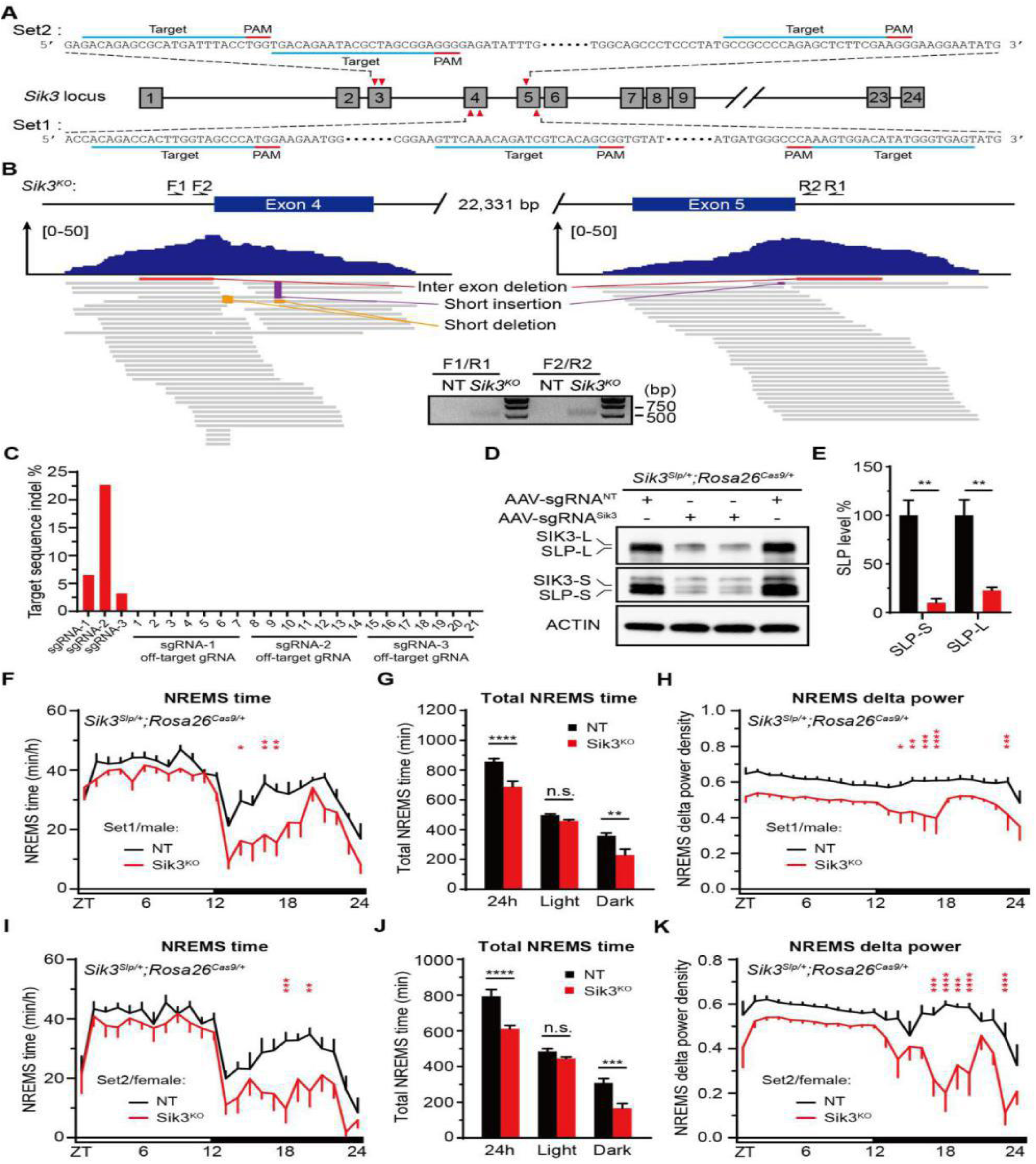
ABC-CRISPR of *Slp/Sik3* rescues hypersomnia of *Sik3^Slp/+^;Rosa26^Cas9/+^* mice. **(A)** Schematic showing two sets of triple sgRNA target sites within exons 3, 4 and 5 of *Sik3* gene. **(B)** Genomic alignments of whole genome sequencing reads of ABC-*Sik3^KO^* mouse brain around the sgRNA target sites within exons 4 and 5 of *Sik3* gene. Different types of mutations are highlighted based on the whole genome sequencing data. Inset shows genomic PCR result with two primer pairs to detect the large inter-exon deletions. **(C)** Quantitative analysis of indel mutations at the sgRNA target sites and twenty-one predicted off-target sites for the three sgRNAs (set 1) targeting *Sik3* gene. **(D)** Immunoblotting of whole brain lysates from AAV-sgRNA^NT^ and AAV-sgRNA*^Sik3^* (set 1) injected *Sik3^Slp/+^; Rosa26^Cas9/+^* male mice with anti-SIK3 and anti-ACTIN antibodies. **(E)** Quantification of the levels of SIK3-L/SLP-L and SIK3-S/SLP-S proteins shown in (D) (n=3). **(F-H)** Hourly plot of NREMS time (F), quantification of total NREMS time (G) and hourly plot of NREMS delta power (H) in the AAV-sgRNA^NT^ (n=7) or AAV-sgRNA*^Sik3^* (set 1, n=7) injected *Sik3^Slp/+^*; *Rosa26^Cas9/+^* male mice. **(I-K)** Hourly plot of NREMS time (I), quantification of total NREMS time (J) and hourly plot of NREMS delta power (K) in the AAV-sgRNA^NT^ (n=7) or AAV-gRNA*^Sik3^* (set 2, n=7) injected *Sik3^Slp/+^*; *Rosa26^Cas9/+^* female mice. Data are mean ± s.e.m. (E) Unpaired t test. (F-K) Two-way ANOVA with Dunn’s multiple comparisons test. n.s. not significant; * *p* < 0.05; ** *p* < 0.01; *** *p* < 0.001; **** *p* < 0.0001.

**Figure 6-figure supplement 1.**
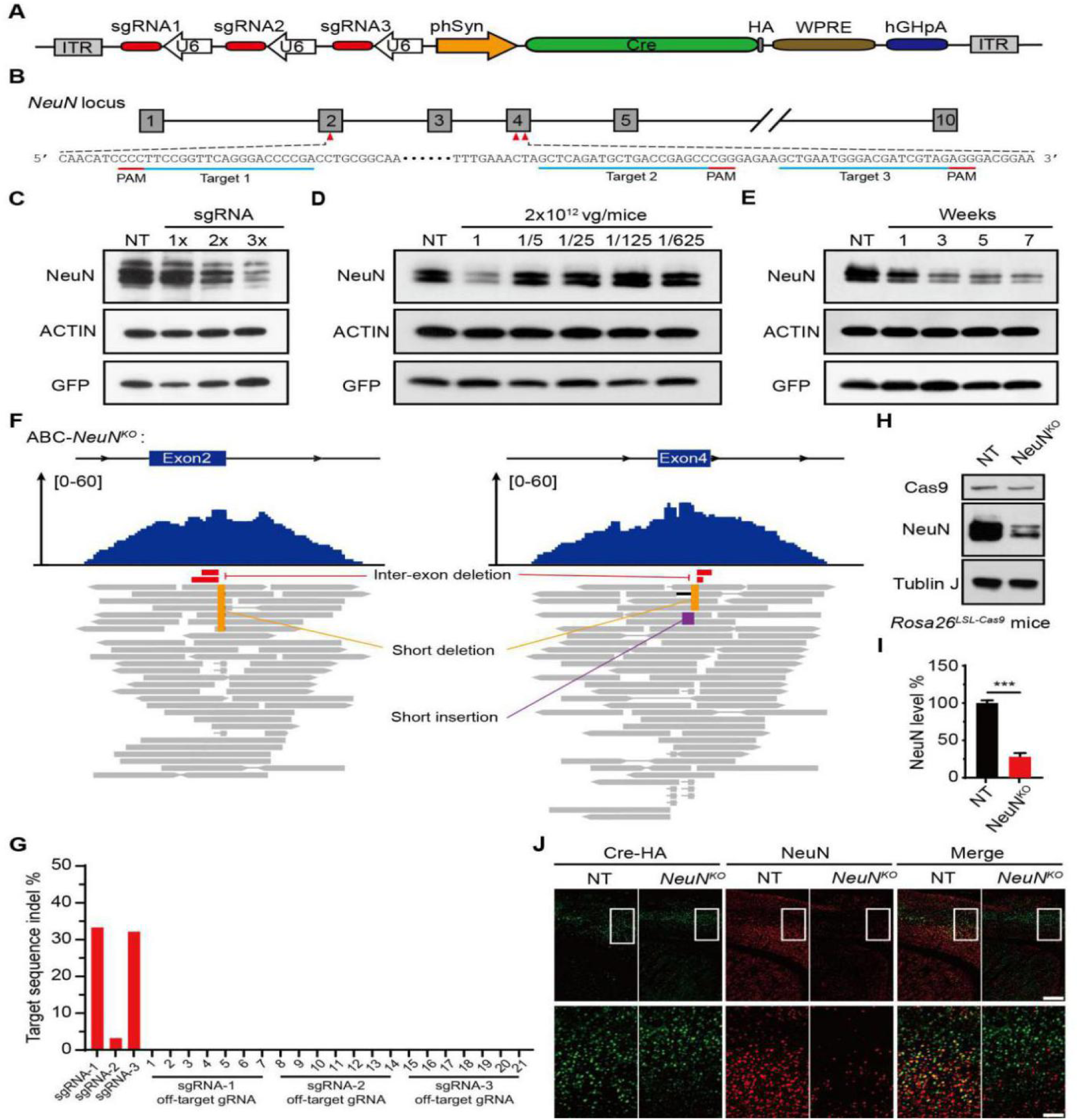
ABC-KO of target genes by triple-target CRISPR in Cas9 mice. **(A)** Schematic of AAV-3xsgRNA*^NeuN^* that expresses HA-Cre recombinase from hSyn promoter and three U6:sgRNA cistrons. **(B)** Schematic of target site sequences within exons 2 and 4 of *NeuN* gene. Immunoblotting of NeuN proteins in brain lysates from AAV-3xsgRNA^NT^, AAV-1xsgRNA*^NeuN^*, AAV-2xsgRNA*^NeuN^*, AAV-3xsgRNA*^NeuN^* (10^12^ vg/mice) injected *Rosa26^LSL-Cas9^* mice. **(D)** Immunoblotting of NeuN proteins in brain lysates from Cas9 mice injected with different doses of AAV-3xsgRNA*^NeuN^*. **(E)** Immunoblotting of NeuN proteins in brain lysates from AAV-3xsgRNA*^NeuN^* injected Cas9 mice at one, three, five or seven weeks after (10^12^ vg/mice) virus injection. AAV-3xsgRNA^NT^ injected mouse brains were collected at three weeks after virus injection. **(F)** Genomic alignments of whole genome sequencing reads of ABC-*NeuN^KO^* mouse brain DNA at the target sites within exons 2 and 4 of *NeuN* gene. The top and bottom panels show read coverage and read alignments, respectively, with different types of mutations highlighted. **(G)** Quantitation of indel mutations at the three target sites and twenty-one predicted off-target sites for the three sgRNAs targeting *NeuN* gene based on the whole genome sequencing data. **(H)** Immunoblotting of whole brain lysates from AAV-3xsgRNA^NT^ and AAV-3xsgRNA*^NeuN^* injected *Rosa26^LSL-Cas9^* mice with the corresponding antibodies. **(I)** Quantification of the level of NeuN expression in (H) (n=4). **(J)** Co-immunostaining of HA-Cre and NeuN in the prefrontal cortex sections of AAV-3xsgRNA^NT^ and AAV-3xsgRNA*^NeuN^* injected mice. The bottom row shows magnified images of the corresponding boxed regions in the top row. Scale bars, 400 μm (top) and 100 μm (bottom). Data are mean±s.e.m. (I) Unpaired t test. n.s. not significant; * *p*<0.05; ** *p*<0.01; *** *p*<0.001; **** *p*<0.0001.

**Figure 6-figure supplement 2.**
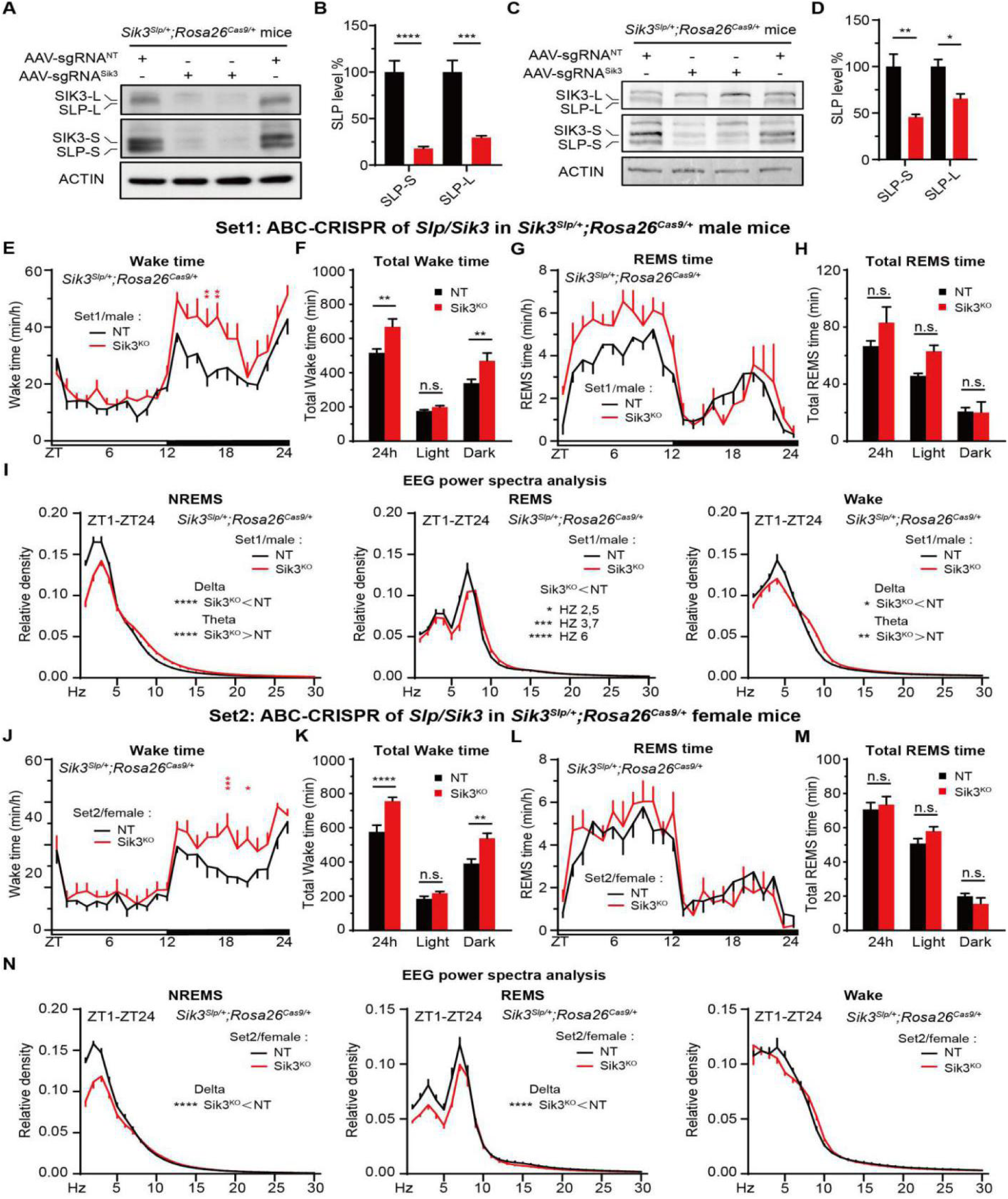
ABC-CRISPR of *Slp/Sik3* rescues hypersomnia of *Sik3^Slp/+^;Rosa26^Cas9/+^* mice. **(A)** Western blotting (ECL) of whole brain lysates from AAV-sgRNA^NT^ and AAV-sgRNA*^Sik3^* (set 2) injected *Sik3^Slp/+^;Rosa26^Cas9/+^* female mice with anti-SIK3 and anti-ACTIN antibodies. **(B)** Quantification of the levels of SIK3-L/SLP-L and SIK3-S/SLP-S proteins shown in (A) (n=4). **(C)** Alkaline phosphatase-based linear Western blotting of the same membrane used in (A) with anti-SIK3 and anti-ACTIN antibodies. **(D)** Quantification of the levels of SIK3-L/SLP-L and SIK3-S/SLP-S proteins shown in (C) (n=4). **(E-I)** Hourly plots of Wake (E) or REMS (G) time, quantification of total Wake (F) or REMS (H) time and EEG power spectra analysis of NREMS, REMS and Wake states (I) in the AAV-sgRNA^NT^ (n=7) or AAV-sgRNA*^Sik3^* (set 1, n=7) injected *Sik3^Slp/+^*; *Rosa26^Cas9/+^* male mice. **(J-N)** Hourly plots of Wake (J) or REMS (L) time, quantification of total Wake (K) or REMS (M) time and EEG power spectra analysis of NREMS, REMS and Wake states (N) in the AAV-sgRNA^NT^ (n=6) or AAV-sgRNA*^Sik3^* (set 2, n=7) injected *Sik3^Slp/+^*;*Rosa26^Cas9/+^* female mice. Data are mean ± s.e.m. (B and D) Unpaired t test. (E-N) Two-way ANOVA with Dunn’s multiple comparisons test. n.s. not significant; * *p* < 0.05; ** *p* < 0.01; *** *p* < 0.001; **** *p* < 0.0001.

### Multiplex ABC-CRISPR of orexin/hypocretin receptors causes narcolepsy-like episodes

The deficiency of neuropeptide orexin/hypocretin or its receptors, OX1R/HCRTR1 and OX2R/HCRTR2 (hereafter OX1R and OX2R for simplicity) results in narcolepsy-like phenotypes, such as abnormal wake to REMS transition and cataplexy, in mice (Chemelli et al., 1999; Kalogiannis et al., 2010; Kohlmeier et al., 2013). For double ABC-CRISPR of OX1R and OX2R, we co-injected Cas9-expressing mice with two AAV-PHP.eB viruses expressing separate sets of triple sgRNAs targeting either *Ox1r* or *Ox2r* gene, respectively **(Figure 7A)**. As shown by reverse transcription (RT)-quantitative (q)PCR, the levels of *Ox1r* and *Ox2r* transcripts were reduced by ∼75% and ∼60%, respectively, in the AAV-sgRNA*^Ox1r/Ox2r^* injected (ABC-*Ox1r/Ox2r^DKO^*) relative to AAV-sgRNA^NT^ injected (ABC-NT) mouse brains **(Figure 7B)**. At first, all of the ABC-*Ox1r/Ox2r^DKO^* mice exhibited no sleep abnormality as compared to the ABC-NT control mice. After feeding with chocolates–a known stimulant of narcolepsy (Oishi et al., 2013), three out of eight ABC-*Ox1r/Ox2r^DKO^* mice exhibited frequent narcolepsy-like episodes, which was characterized by the abnormal transitions from wake to REMS during EEG/EMG recording **(Figure 7C-E)**. However, we did not observe cataplexy during the narcolepsy episodes through simultaneous video/EEG recording and manual inspection. This partial narcoleptic phenotype was probably due to incomplete knockout of OX1R and OX2R in the adult brain neurons. These results suggest that multiplex ABC-CRISPR can enable one-step analysis of the sleep phenotypes of redundant genes in adult mice.

**Figure 7.**
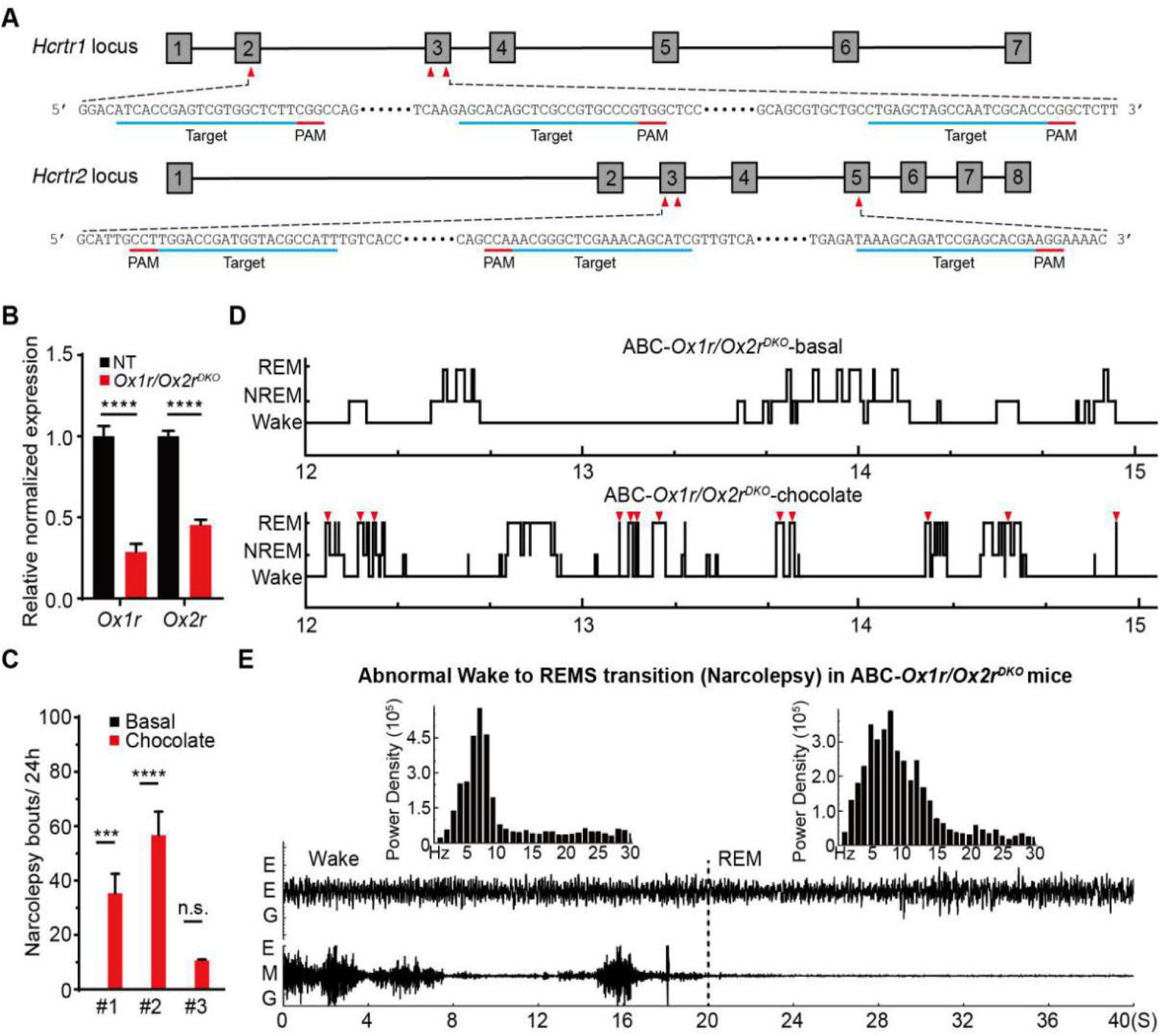
Multiplex ABC-CRISPR of *Ox1r* and *Ox2r* causes chocolate-induced narcolepsy episodes. **(A)** Schematic showing the two sets of sgRNA target sites within exons 2 and 3 of *Ox1r* gene and exons 3 and 5 of *Ox2r* gene. **(B)** Quantification of *Ox1r* and *Ox2r* transcript levels in the ABC-NT and ABC-*Ox1r/Ox2r^DKO^* mouse brains by real-time reverse-transcription (RT)-quantitative PCR. **(C)** Quantification of the number of narcolepsy episodes in three ABC-*Ox1r/Ox2r^DKO^* mice before and after chocolate feeding for three days. **(D)** Representative hypnograms (ZT12-15) of ABC-*Ox1r/Ox2r^DKO^* mice before and after chocolate feeding. Red triangles mark abnormal wake to REM transitions during narcolepsy-like episodes. **(E)** Representative EEG/EMG signals depicting the abnormal Wake to REM transition during one narcoleptic episode in the ABC- *Ox1r/Ox2r^DKO^* mice. Data are mean ± s.e.m. (B and C) Unpaired t test. n.s. not significant; * *p* < 0.05; ** *p* < 0.01; *** *p* < 0.001; **** *p* < 0.0001.

### ABC-expression/KO screens identify Ankrd63 and NR1 as potentially new sleep regulators

We recently investigated the molecular substrates of homeostatic sleep need by comparing whole brain phosphoproteomes of *Sleepy* mutant and sleep-deprived wild-type mice (Wang et al., 2018). We identified 80 sleep need index phosphoproteins (SNIPPs), whose phosphorylation states change in relation to sleep need in these two opposite models of high sleep need. The majority of SNIPPs are annotated synaptic proteins, implicating a potential mechanistic link between synaptic phosphoproteome and sleep need regulation. Furthermore, mutant SLP kinase preferentially associates with SNIPPs, suggesting that they may function as downstream substrates of SLP kinase and effectors of homeostatic sleep regulation.

To identify new sleep regulators, we performed ABC-expression screening of eight candidate SNIPPs, including Abl-interactor-1(Abi1) (Rial Verde et al., 2006), Ankyrin repeat and sterile alpha motif domain-containing protein 1B (Anks1b) (Tindi et al., 2015), ADP-ribosylation factor-GTPase-activating proteins 2 (Arfgap2) (Saitoh et al., 2009), ADP-ribosylation factor-GTPase-activating proteins 3 (Arfgap3) (Dembla et al., 2014), Ankyrin repeat domain-containing protein 63 (Ankrd63), Synaptopodin (Synpo) (Zhang et al., 2013), DEAD-Box Helicase 3 Y-linked (Ddx3y) (Vakilian et al., 2015), Pyruvate dehydrogenase E1 subunit alpha 1 (Pdha1) (Jakkamsetti et al., 2019), and followed by EEG/EMG-based sleep analysis. As shown by immunoblotting **(Figure 8A)**, seven of eight SNIPPs were expressed at substantial levels in the adult mouse brains following intravenous AAV injection. Among these, only ABC-expression of Ankrd63 resulted in significant sleep phenotypes relative to ABC-eGFP and other ABC-SNIPP mice **(Figure 8B)**. Specifically, ABC-Ankrd63 mice showed reduction in NREMS time and NREMS delta power, increase of wake time and no change in REMS time **(Figure 8C-E and Figure 8-figure supplement 1A-E)**.

**Figure 8.**
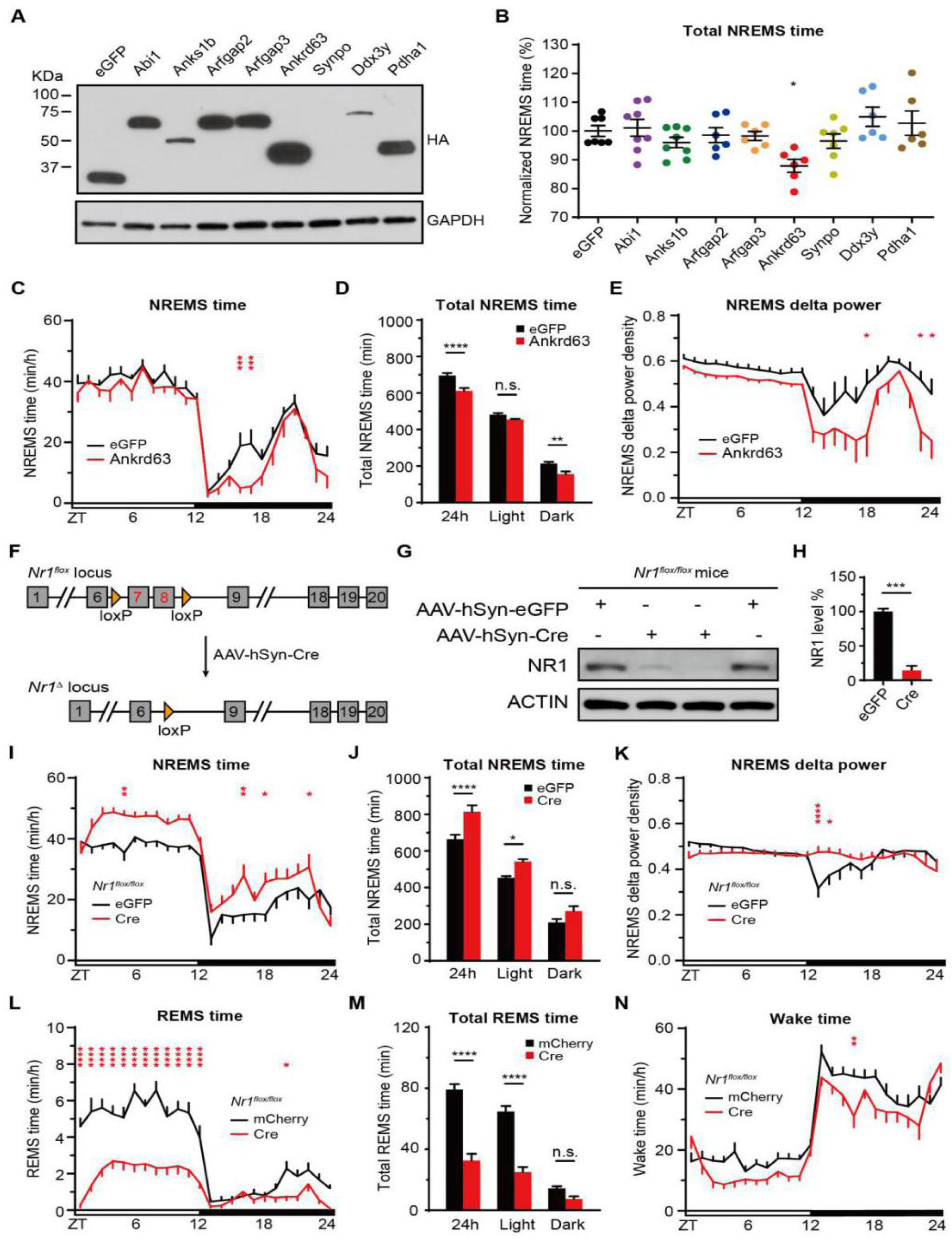
ABC-expression/KO screen identifies Ankrd63 and NR1 as two new sleep regulators. **(A)** Immunoblotting of whole brain lysates from AAV-hSyn-SNIPPs injected C57BL/6J adult mice with anti-HA and anti-GAPDH antibodies. **(B)** Daily NREMS time of ABC-SNIPPs mice (n ≥ 5) normalized to that of ABC-eGFP mice. **(C-E)** Hourly plots of NREMS time (C), quantification of total NREMS time (D) and hourly plot of NREMS delta power (E) in the ABC-eGFP (n=7) or ABC-Ankrd63 (n=6) mice. **(F)** Schematic of ABC-KO of *Nr1* by intravenous injection of AAV-hSyn-Cre in *Nr1^flox/flox^* mice. **(G)** Immunoblotting of brain lysates from AAV-hSyn-eGFP or AAV- hSyn-Cre injected *Nr1*^flox/flox^ mice with anti-NR1 and anti-ACTIN antibodies. **(H)** Quantification of the level of NR1 proteins in (G) (n=3). **(I-N)** Hourly plots of NREMS (I), REMS (L) or Wake (N) time, quantification of total NREMS (J) or REMS (M) time and hourly plot of NREMS delta power (K) in the AAV-hSyn-eGFP (n=15) or AAV-hSyn-Cre (n=15) injected *Nr1^flox/flox^* mice. Data are mean ± s.e.m. (B) One-way ANOVA with Dunn’s multiple comparisons test. (C-E and I-N) Two-way ANOVA with Dunn’s multiple comparisons test. (H) Unpaired t test. n.s. not significant; * *p* < 0.05; ** *p* < 0.01; **** *p* < 0.0001.

Previously, Ueda and colleagues have reported that *Nr3a* KO mice exhibit a short sleep phenotype by using triple-target CRISPR to generate whole-body biallelic KO mice for members of the N-methyl-D-aspartic (NMDA) receptor family (Sunagawa et al., 2016). However, the sleep functions of essential genes *Nr1* and *Nr2b* could not be analyzed owning to embryonic lethality of KO mice. Here, we performed ABC-KO of *Nr1* by injection of AAV-hSyn-Cre into *Nr1^flox/flox^* adult mice, in which exons 7 and 8 of *Nr1* gene were flanked by two loxP sites **(Figure 8F)**. Immunoblotting confirmed efficient KO of NR1 expression in the majority of adult brain neurons **(Figure 8G and 8H)**. Accordingly, ABC-KO of *Nr1* caused on average ∼150 min increase in daily NREMS amount, ∼47 min (∼60%) decrease in daily REMS amount, and ∼100 min decrease in total wake time **(Figure 8I-N and Figure 8-figure supplement 1F)**. It is worth noting that *Nr3a^KO^* and ABC-*Nr1^KO^* mice showed opposite changes in NREMS amount, suggest a complex role of NMDA receptor in the regulation of daily sleep time. Taken together, this candidate ABC-expression/KO screen identified Ankrd63 and NR1 as two potentially new sleep regulators.

**Figure 8-figure supplement 1.**
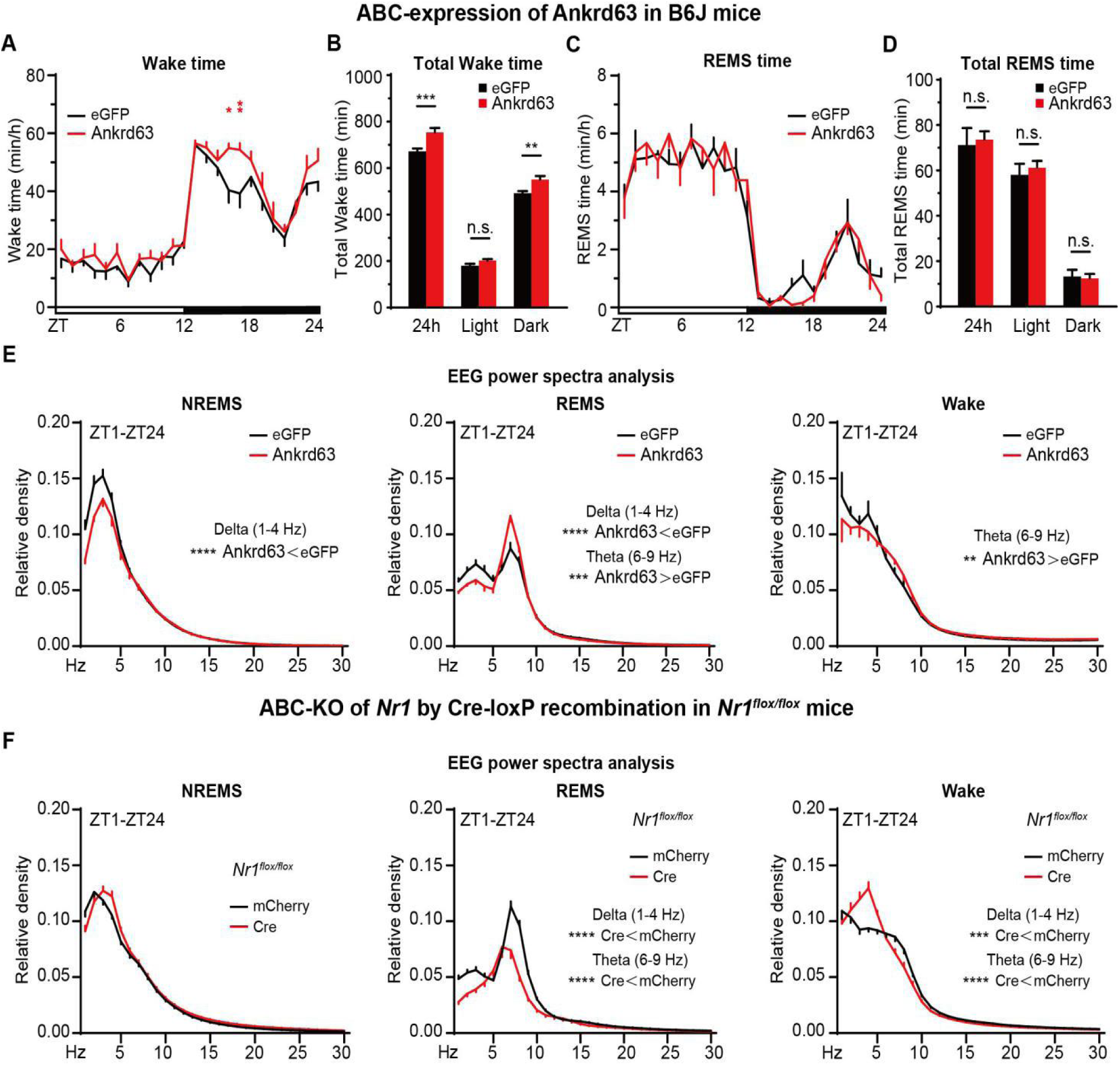
ABC-expression/KO screen identifies Ankrd63 and NR1 as two new sleep regulators. **(A-E)** Hourly plots of Wake (A) or REMS (C) time, quantification of total Wake (B) or REMS (D) time and EEG power spectra analysis of NREMS, REMS and Wake states (E) in the ABC-eGFP (n=7) or ABC-Ankrd63 (n=6) injected mice. **(F)** EEG power spectra analysis of NREMS, REMS and Wake states in the AAV-hSyn-eGFP (n=15) or AAV-hSyn-Cre (n=15) injected *Nr1^flox/flox^* mice. Data are mean ± s.e.m. (A-F) Two-way ANOVA with Dunn’s multiple comparisons test. n.s. not significant; * *p* < 0.05; ** *p* < 0.01; *** *p* < 0.001; **** *p* < 0.0001.

## DISCUSSION

### Rapid and efficient somatic genetics analysis of sleep in adult mice

The molecular mechansims of mammalian sleep regulation remain largely unknown. Forward genetics screening of randomly mutagenized mice is a powerful and hypothesis-free approach to identify key sleep regulatory genes in mammals (Banks et al., 2020; Funato et al., 2016; Kapfhamer et al., 2002; Takahashi et al., 1994). On the other hand, reverse genetics, through the making of transgenic, knock-in, KO and conditional KO mice for specific genes of interest, represents a hypothesis-driven approach to identify and characterize new sleep regulatory genes (Graves et al., 2003; Hellman et al., 2010; Honda et al., 2018; Mikhail et al., 2017; Takahashi et al., 1994). This latter approach is greatly accelerated by next generation gene-editing technologies, such as the CRISPR/Cas9 system (Hsu et al., 2014; Sunagawa et al., 2016; Tatsuki et al., 2016; Wang et al., 2013). However, classical forward and reverse genetics approaches involve germline mutations, therefore, are unwieldy to study the essential genes (owning to early lethality) or redundant genes (owning to lack of phenotype) **(Table 1)**. In some but not all cases, forward or reversed genetics may identify viable gain-of-function or partial loss-of-function mutations that can uncover the sleep phenotypes of essential genes (Funato et al., 2016; Graves et al., 2003).

**Table 1.**
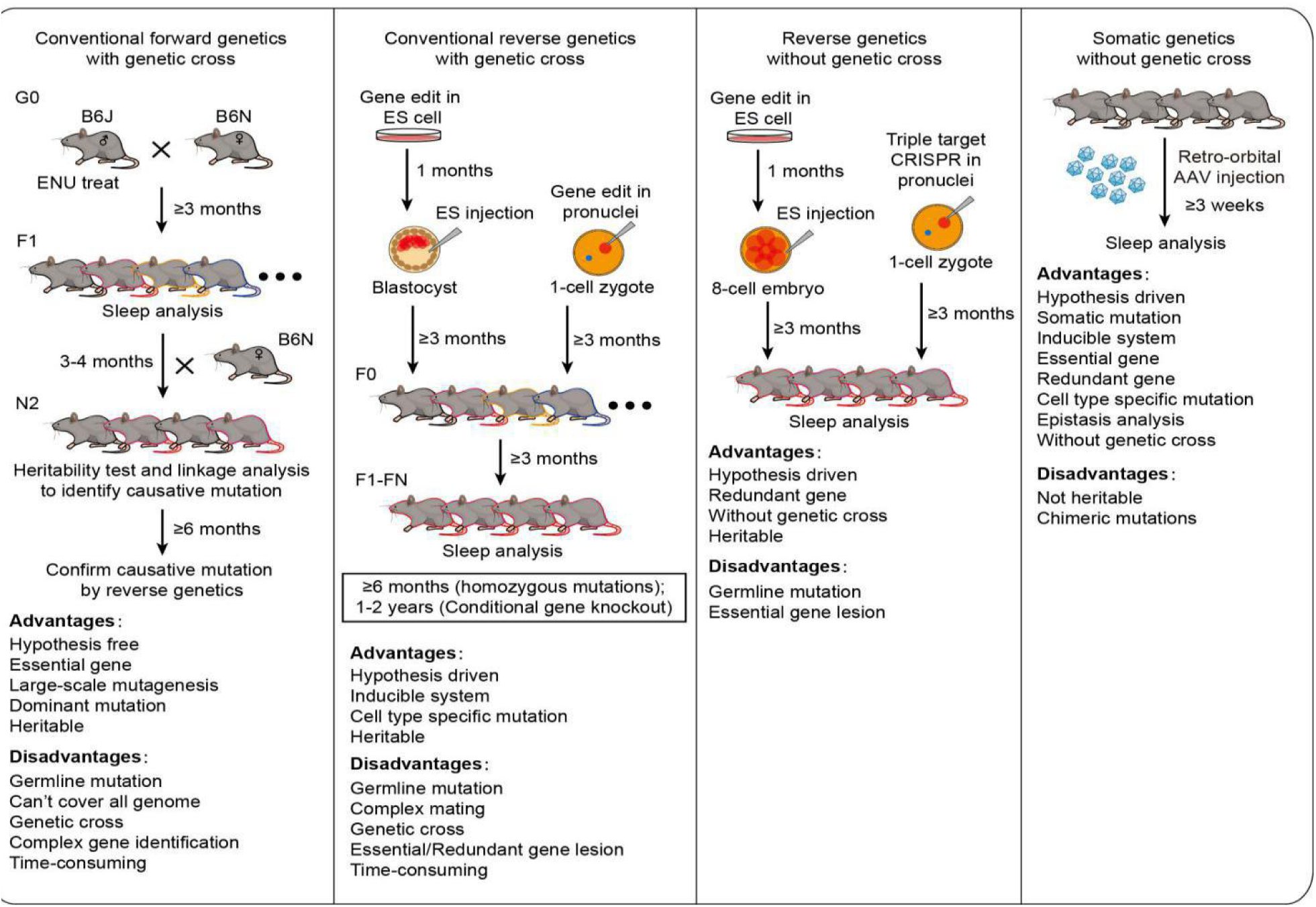
Comparison between forward/reverse genetics and somatic genetics approaches in mouse sleep study.

In this study, we applied the AAV-based ABC-expression/KO platform for somatic genetics analysis of sleep in adult mice **(Table 1)**. We conducted two pilot ABC sleep screens that identified CREB and three potentially new sleep regulators: CRTC1, Ankrd63 and NR1. Moreover, both constitutive and Tet-inducible expression of CREB and CRTC1 significantly reduced daily NREMS amount and NREMS delta power. There are two well-known mechanisms for the activation of CREB activity: a) phosphorylation of S133 of CREB; b) CRTCs function as co-activators for CREB. Our results suggest that the ability of CREB to regulate sleep quantity/quality is probably independent of S133 phosphorylation, but can be enhanced by CRTC1. These results demonstrate the proof-of-principle that this somatic genetics approach should facilitate rapid identification of new sleep regulatory genes, especially from essential genes, in adult mice by skipping the mouse development.

Moreover, the ABC-expression/KO platform is a highly efficient approach to conduct sophisticated genetic experiments in adult mice without genetic crosses. For example, when ABC-expression or KO of two different genes produce opposite sleep phenotypes, epistasis analysis can be rapidly carried out to determine whether the two genes operate in the same or parallel pathways and map the order of these genes if they are in the same pathway. For structural and functional analysis of key sleep regulators, ABC-expression of wild-type and mutant proteins can be conducted to rescue the sleep phenotypes of ABC-KO of endogenous proteins. Furthermore, a suppressor screen can be easily conducted to identify the downstream effectors of SLP/SIK3 kinases in sleep regulation by ABC-expression of candidate proteins in *Sik3^Slp/+^* mice, or by ABC-CRISPR of target genes in *Sik3^Slp/+^*;*Rosa26^Cas9/+^* mice. Similarly, the ABC-expression/KO platform can be used for enhancer screens to uncover the redundant pathways of sleep regulation. Thus, this somatic genetics approach should also facilitate efficient characterization of the molecular pathways of sleep regulation in mice.

### ABC-KO by AAV-Cre injection facilitates systematic screening of conditional flox mice

To bypass embryonic lethality, conditional flox mice are sometimes crossed with CaMKIIα-Cre transgenic mice, which begin to express Cre recombinase during the third postnatal week (Tsien et al., 1996). Alternatively, conditional flox mice can be crossed with transgenic mice expressing from a pan-neuronal promoter the Cre^ERT2^ fusion proein between Cre and mutant estrogen receptor, which only becomes activated and translocates to the nucleus upon binding of synthetic 4-hydroxytamoxifen (Feil et al., 1996). In theory, the sleep phenotypes could be evaluated in adult mice after tamoxifen injection to induce Cre-dependent KO of target gene in the adult brain neurons. However, both methods have significant drawbacks: 1) CaMKIIα-Cre mice express Cre only in excitatory neurons of the forebrain and Cre expression begins days after birth; 2) Cre^ERT2^- mediated loxP recombination is inefficient after tamoxifen injection in adult mice (Iwasaki et al., 2021).

We showed that intravenous injection of *Creb1^flox/flox^* adult mice with AAV-hSyn-Cre could efficiently knockout CREB expression in the adult brain neurons and significantly increase daily NREMS amount. On the other hand, AAV-hSyn-Cre injection efficiently excised exon 13 of *Sik3* in the adult brain neurons of *Sik3-E13^flox/flox^* mice, which phenocopied germline *Sleepy* mutant mice. Thus, ABC-KO by AAV-mediated Cre/loxP recombination can create either loss- or gain-of-function mutations of target gene for sleep phenotype analysis in the adult mice. It is important to highlight that this simple ABC-KO method is more efficient than traditional method of breeding *synapsin 1-Cre^ERT2^;Sik3-E13^flox/+^* mice, which exhibit little or no sleep phenotype after tamoxifen injection in adult mice (Iwasaki et al., 2021).

A significant recent development for mouse genetics is the availability of a large repertoire of conditional flox strains for most mouse genes from the International Knockout Mouse Consortium (IKMC) and commercial sources. Therefore, we envision that ABC-KO by AAV-Cre injection will facilitate systematic screening of conditional flox mice for sleep and other brain-related phenotypes. Moreover, AAV-delivered Cre expression from various cell type-specific promoters, such as hSyn (neurons), CaMKII (excitatory neurons), mDlx (inhibitory neurons) and etc. (de Leeuw et al., 2016; Dimidschstein et al., 2016; Graybuck et al., 2021; Mich et al., 2021; Tsien et al., 1996), represents a simpler and more efficient method to investigate cell type-specific functions of the target gene in the mouse brain without crossing the flox mice with various Cre-transgenic mice.

### How to distinguish between primary vs. secondary sleep phenotypes?

A large number of genes play important roles in the development of mouse brain. Thus, it is often uncertain whether the sleep phenotype of a mutant mouse strain is the primary phenotype or secondary phenotype owning to developmental abnormalities of the brain. The ABC-expression/knockout platform can be used to effectively address this challenging question by skipping mouse development and directly assessing the sleep phenotypes of somatic mutations in the adult brain neurons. For example, ABC-KO of exon 13 of *Sik3* induced hypersomnia in *Sik3-E13^flox/flox^* mice, whereas ABC-CRISPR of *Slp/Sik3* largely reversed hypersomnia in *Sleepy* (*Sik3^Slp/+^*) mice. These results strongly suggest that the hypersomnia of *Sleepy* mice is the primary phenotype owing to a direct role of mutant SLP kinase in sleep regulation in the adult brain neurons. Thus, this type of somatic genetics analysis can serve as an effective tool to distinguish between the primary and secondary sleep phenotypes for genes that are also important for mouse brain development.

### Multiplex ABC-CRISPR enables one-step analysis of redundant sleep genes

To analyze the phenotypes of redundant genes, it is time-consuming (≥ 2 years) to generate double or triple knockout mice through classical germline genetic approaches (Sunagawa et al., 2016). A combination of triple-target CRISPR and modified embryonic stem cell technologies allows for biallelic knockout of multiple genes in a single generation, however, this strategy only works for non-essential genes (Sunagawa et al., 2016). Here, we successfully used multiplex ABC-CRISPR technology to disrupt both *Ox1r* and *Ox2r* genes in adult mice, resulting in chocolate-induced narcolepsy episodes. It should be noted that the efficiency of multiplex ABC-CRISPR can be further improved by optimizing the sgRNA structure or pre-screening of sgRNAs, and by potentially developing other CRISPR/Cas systems. For example, unlike CRISPR/Cas9 that requires three U6:sgRNA cistrons to target the same gene, CRISPR/Cpf1 (Cas12a) and CRISPR/Cas13d (targeting RNA) can process a polycistronic transcript into multiple gRNAs targeting the same or different genes (Konermann et al., 2018; Zetsche et al., 2017; Zhong et al., 2017). Thus, multiplex ABC-CRISPR will enable one-step analysis of redundant sleep genes in adult mice, which is also applicable for essential genes and can be achieved in one month rather than 1-2 years.

### Concluding statement

We believe that this somatic genetics approach could potentially be useful for studies of many brain-related patho/physiological and behavioral processes. First, it skips embryonic development and directly assess the functions of genes in adult mice without genetic crosses, which is especially useful for studying essential genes or redundant pathways. Second, it allows non-neuroscientists and neuroscientists alike to easily express or ablate target genes in adult moue brains without strong skills in stereotactic injection or prior knowledge of where the genes may function in the mouse brain; Third, it facilitates complex mouse genetic experiments, such as epistasis analysis and enhancer/suppressor screens, with an unprecedented speed that is simply impossible with existing tools. Furthermore, similar strategies can be developed for somatic genetics studies of mouse genes in other peripheral organs by using AAV variants engineered to efficiently transduce the muscle, heart, kidney, intestine, lung, skin, testis and ovary (Chan et al., 2017; Gradinaru, 2020; Pulicherla et al., 2011; Tabebordbar et al., 2021).

## ACKNOWLEDGEMENTS

We are thankful for Drs. M. Luo and F. Shao for sharing reagents; K. Wu, Z. Li, W. Min, Y. Zhuang, H. Huang, and Y. Yin for technical assistance; Drs. X. Wang, F. Shao, R. Xi, and N. Tang for discussion and comments on the manuscript; B. Wu for developing the EEG/EMG analysis software; M. Shi for graphical abstract design. This work was supported by the Natural Science Foundation of China (61772526 to Z.C.), the Beijing Municipal Science and Technology Commission (Z181100001318004 to Q.L.) and the National Key Research and Development Program of China.

## AUTHOR CONTRIBUTIONS

Q.L., G.W., Q.L., J.X., S.Z., R.Z. designed and executed the experiments with help from F.W., Z.C.; G.W. Q.L., J.X., R.Z. developed the ABC-expression/KO platform and performed molecular genetics experiments with help from Z.C., W.J.; S.Z., Q.L., Z.C. developed the SleepV (video) system; X.G., S.Z., Z.C., C.M., L.C., B.S. performed EEG/EMG recording and analysis; F.W., R.Z. generated multiple mouse strains; Q.S., Y.G. performed AAV packaging and purification; H.W., X.W., H.L. helped mouse husbandry. G.W., Q.L., J.X., S.Z., R.Z.prepared the figures and Q.L., wrote the manuscript with help from G.W., Q.L.

## COMPETING INTERESTS

We declare no competing interests.

## Materials and methods

### Key resources table

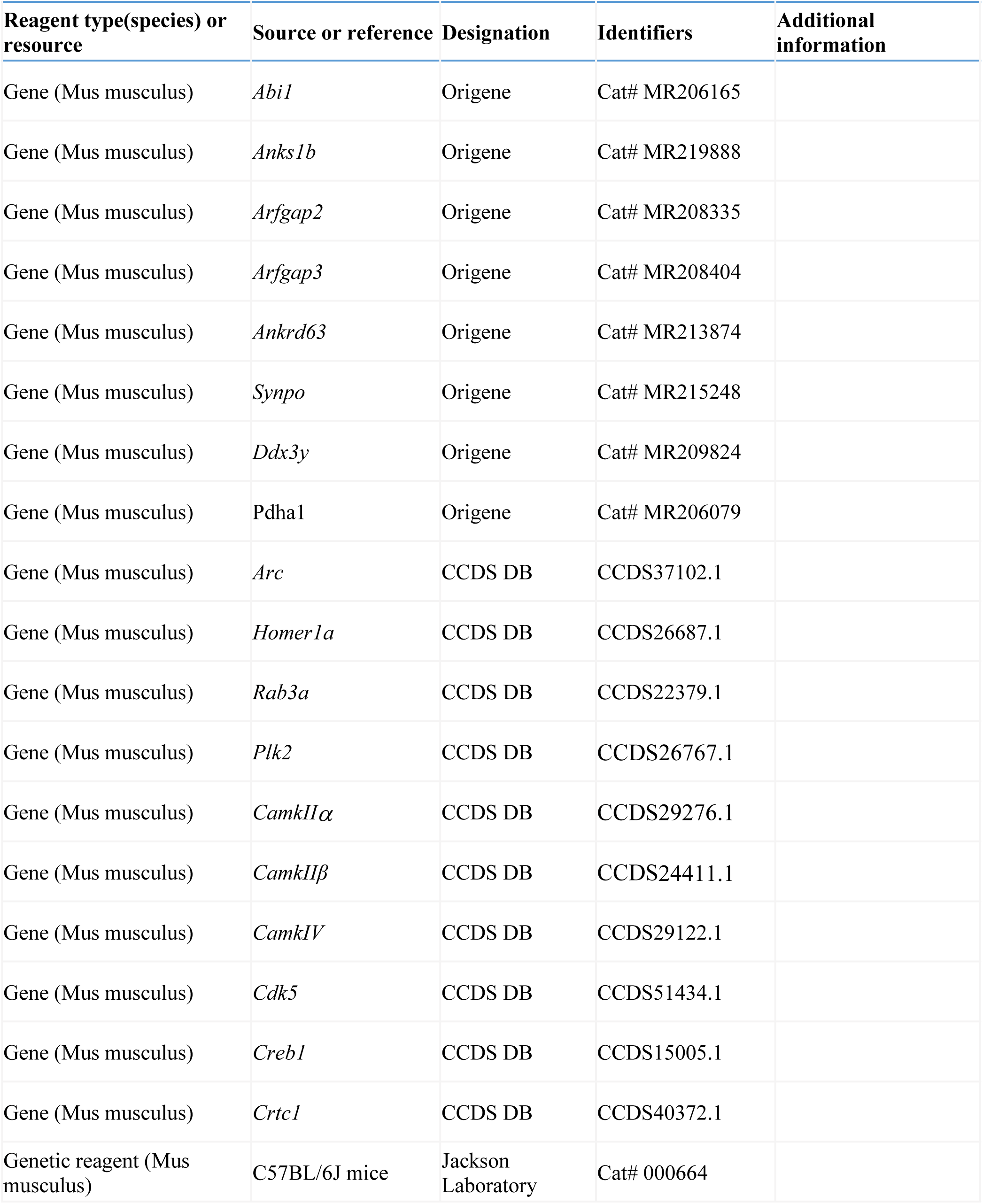
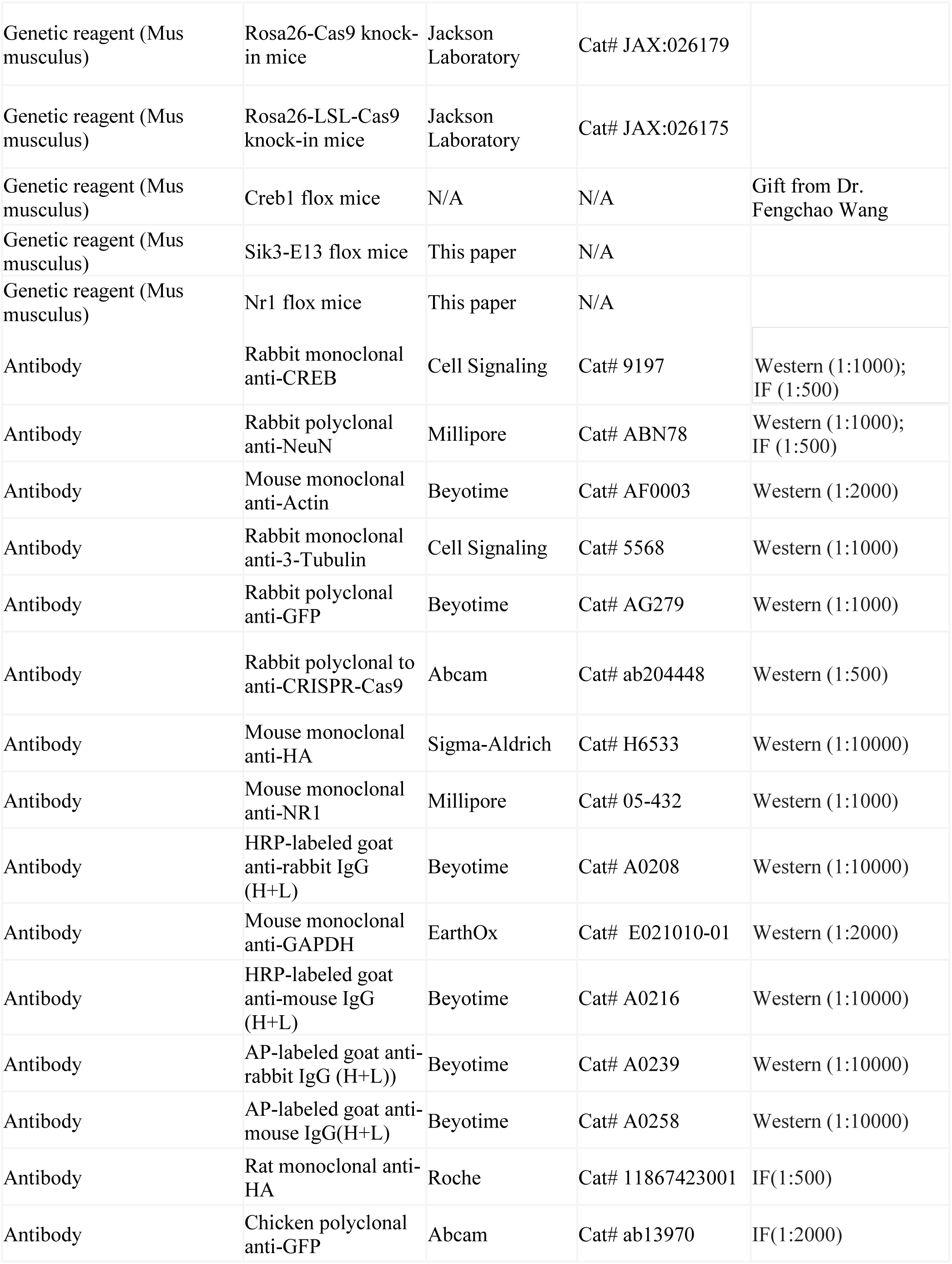
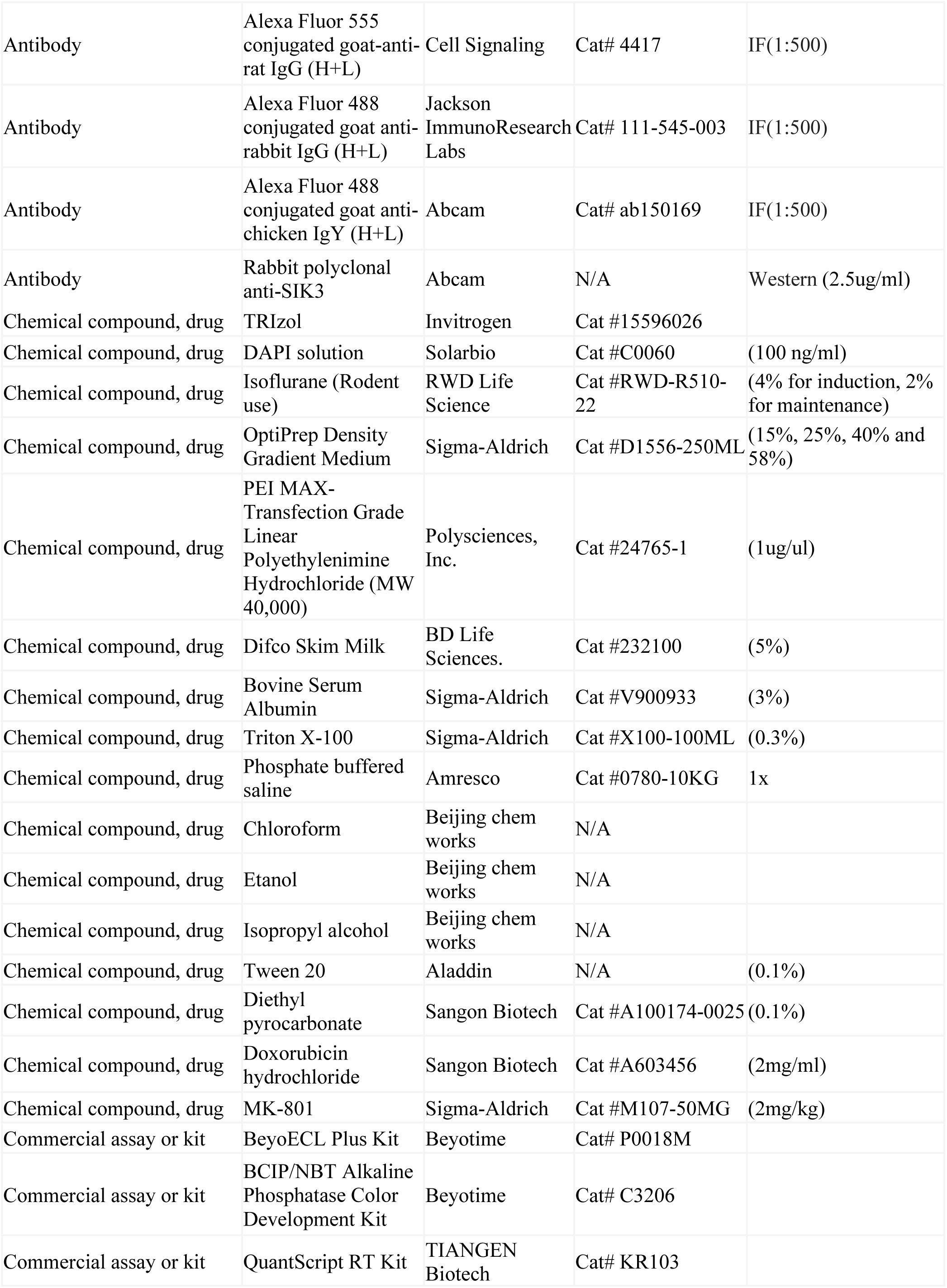
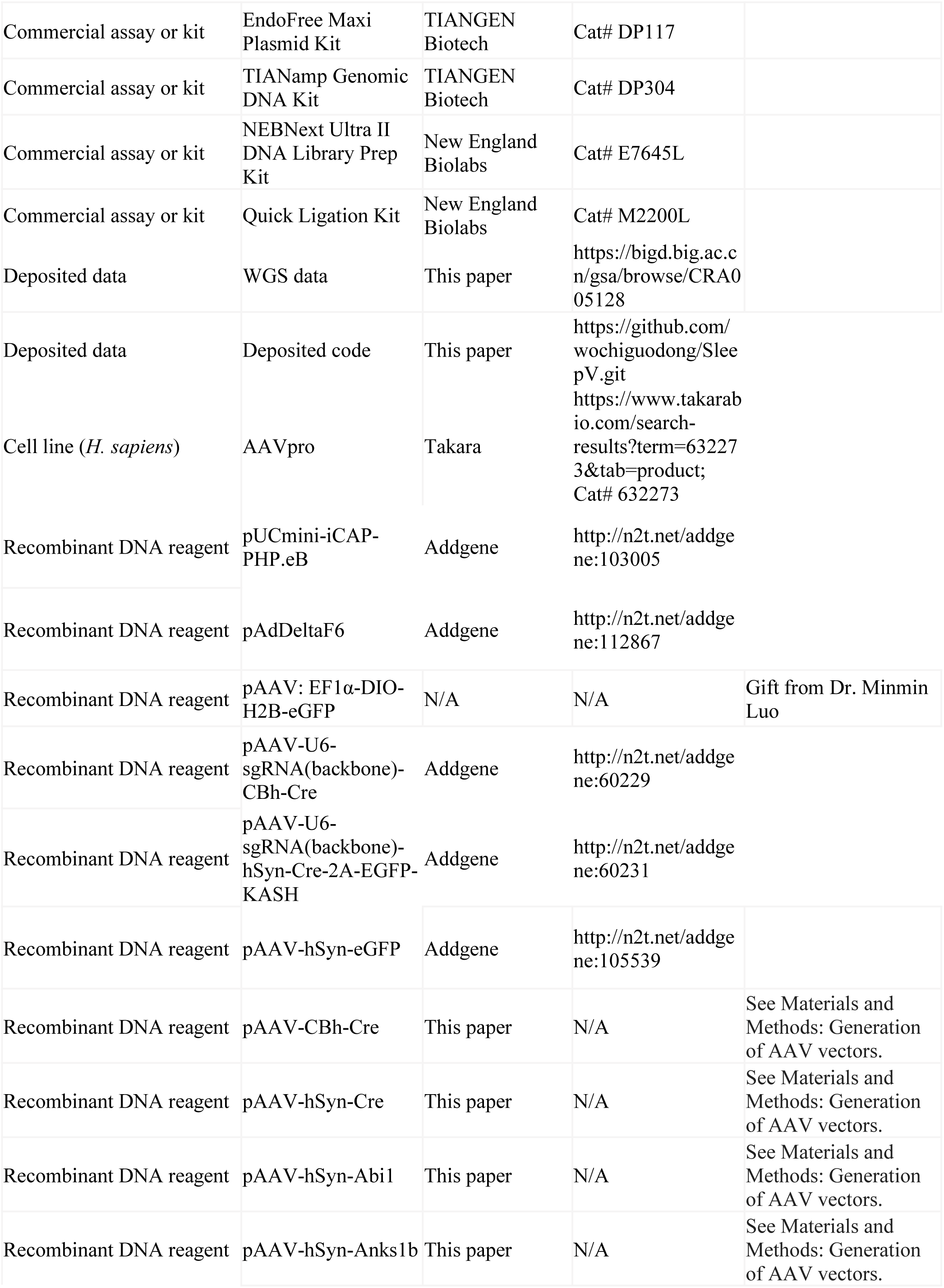
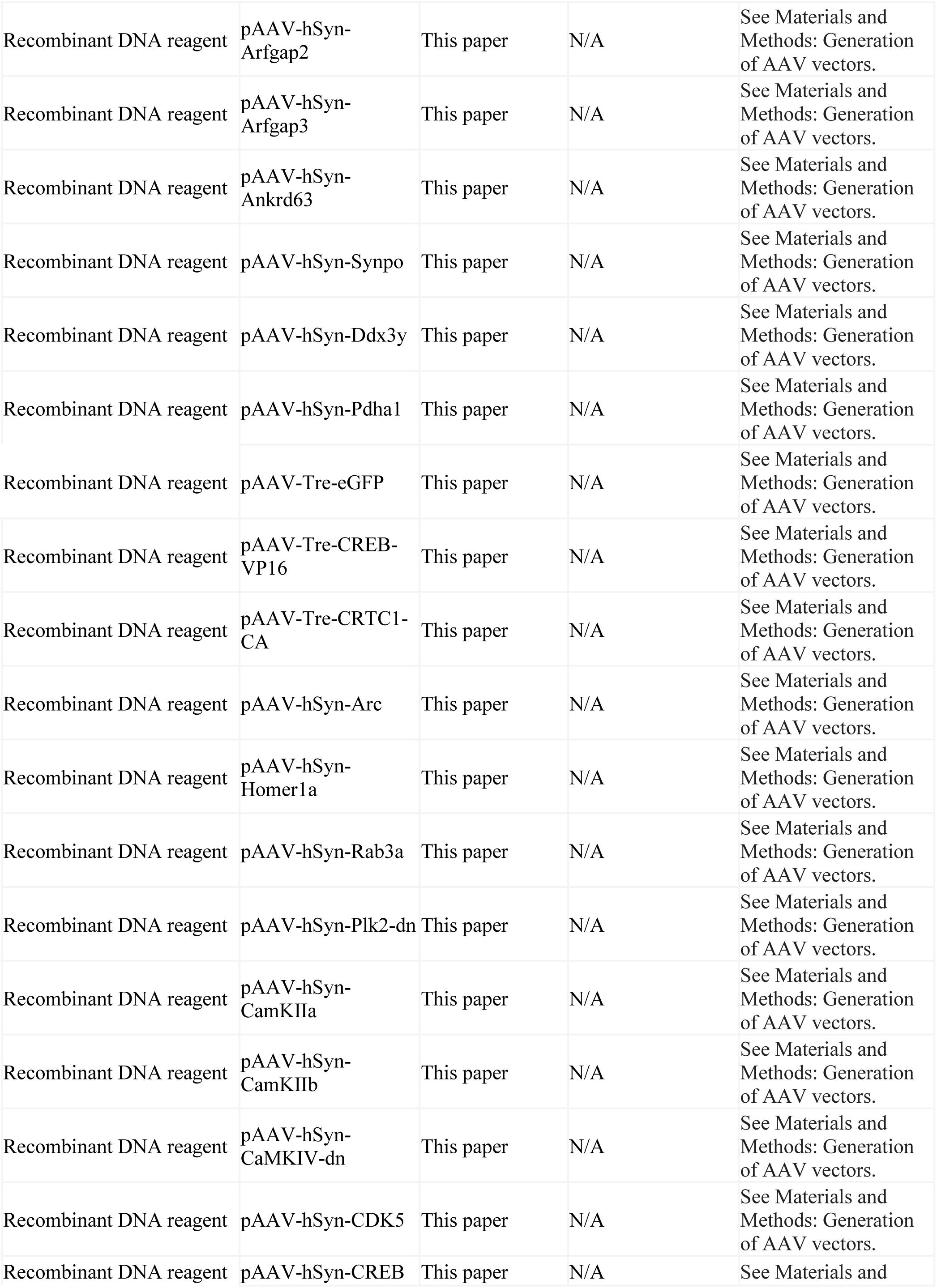
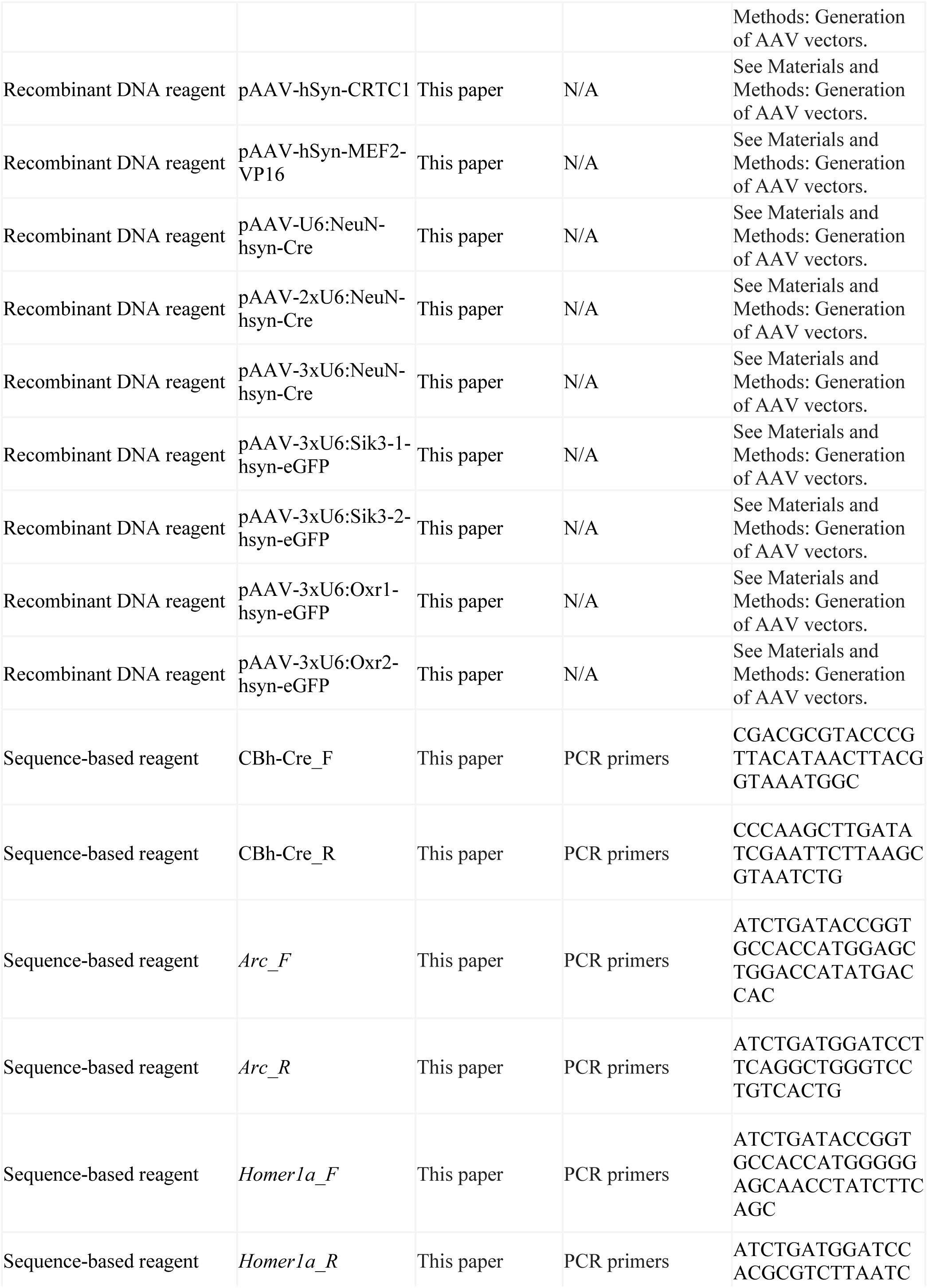
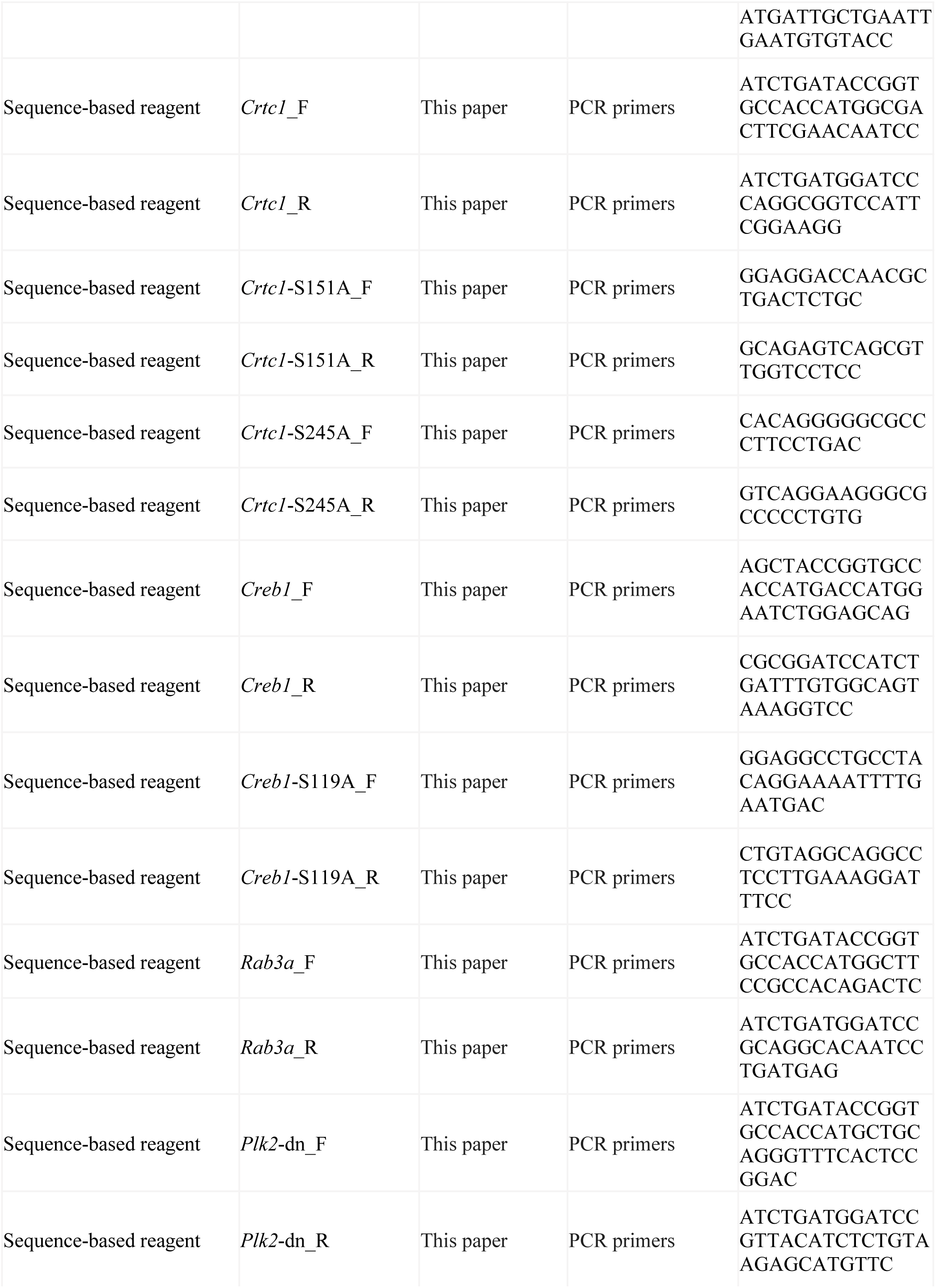
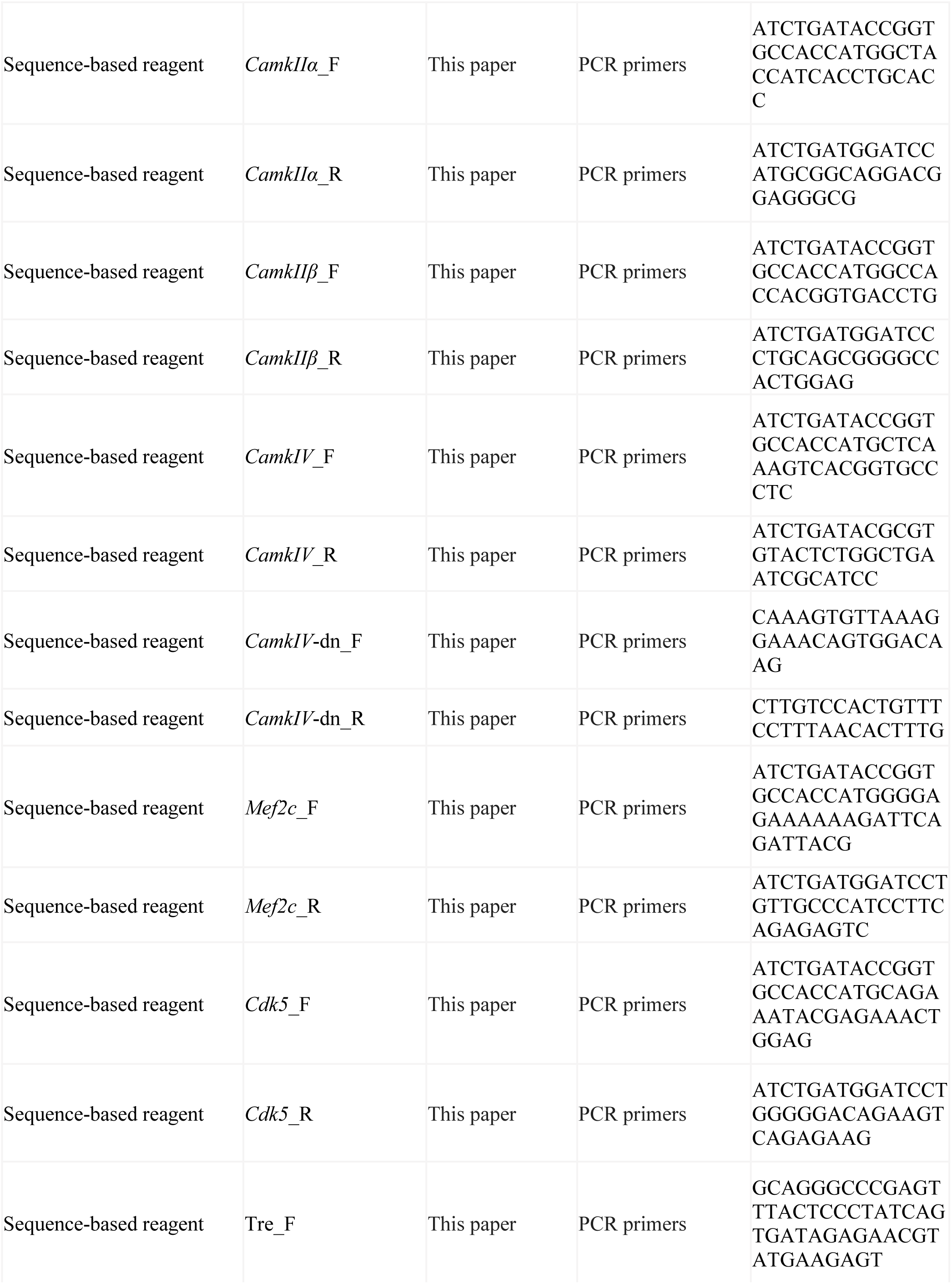
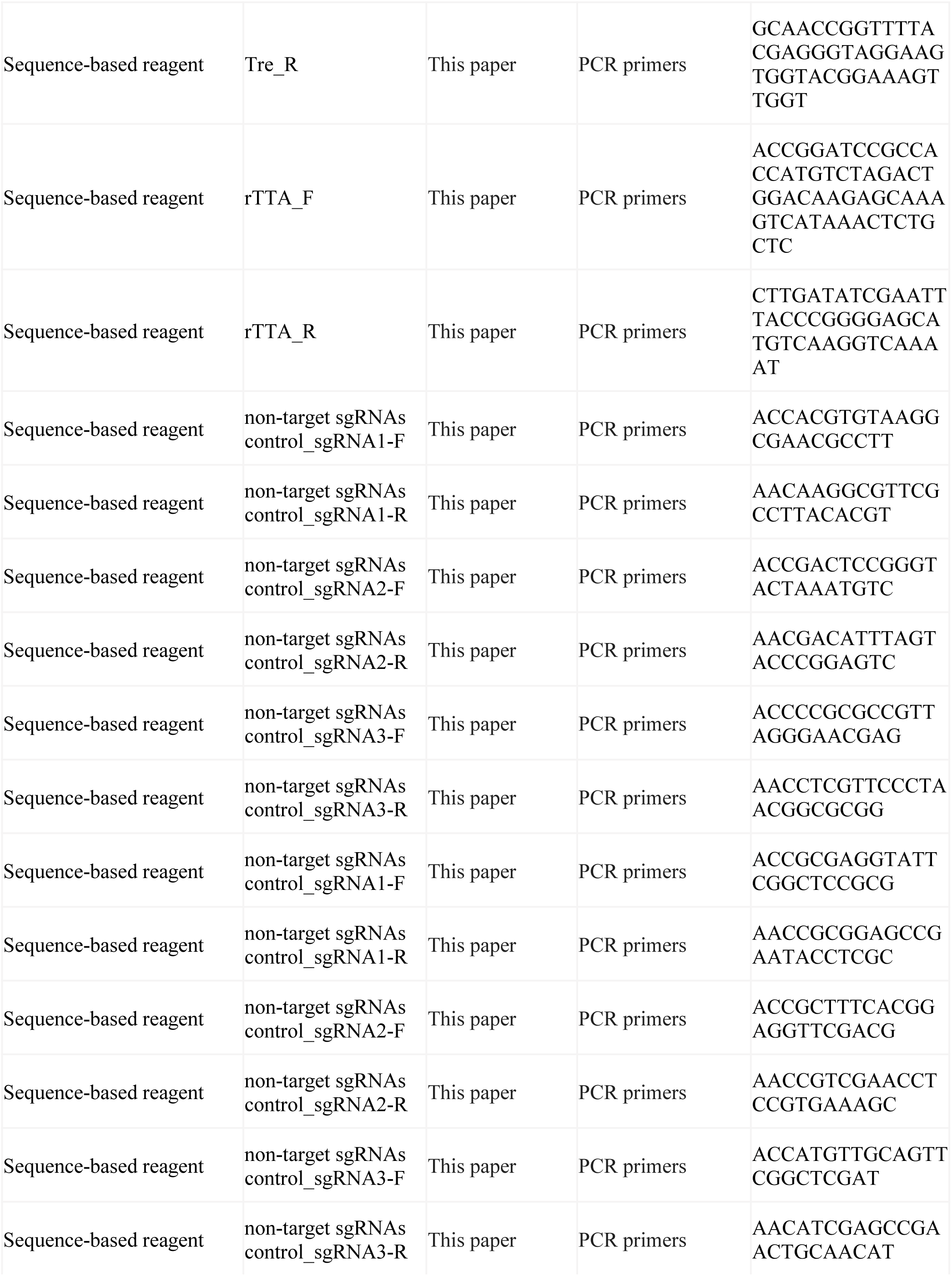
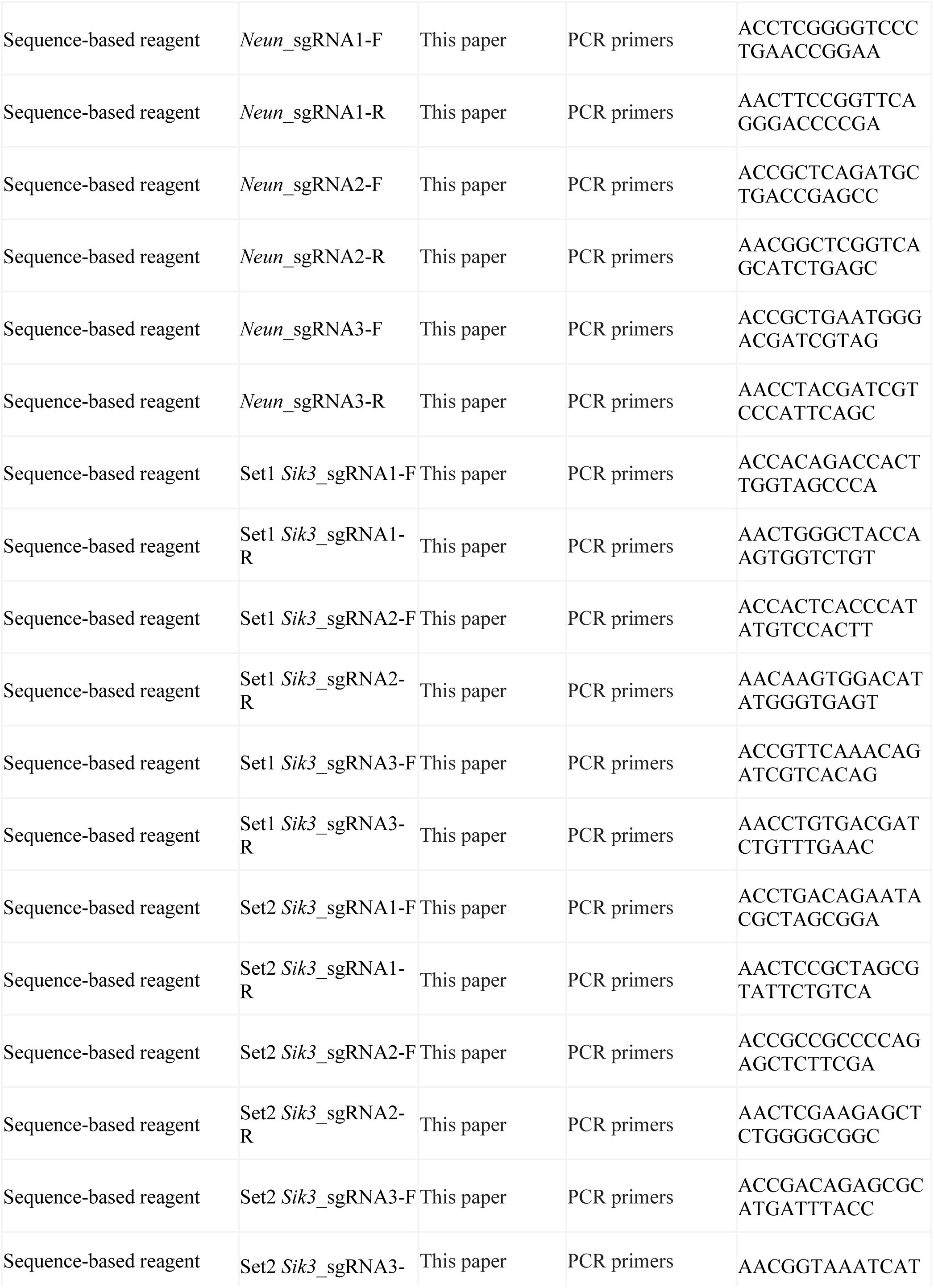
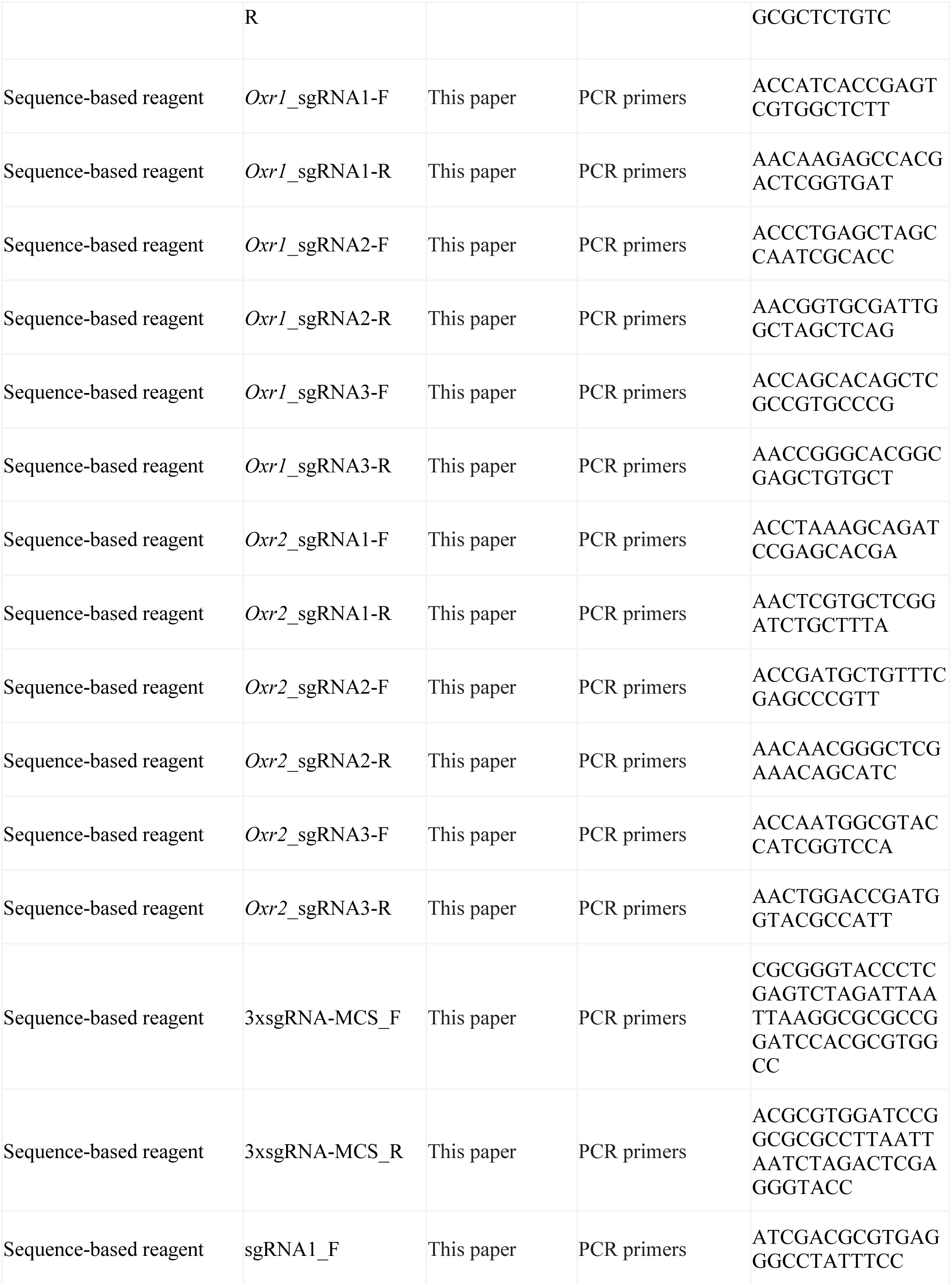
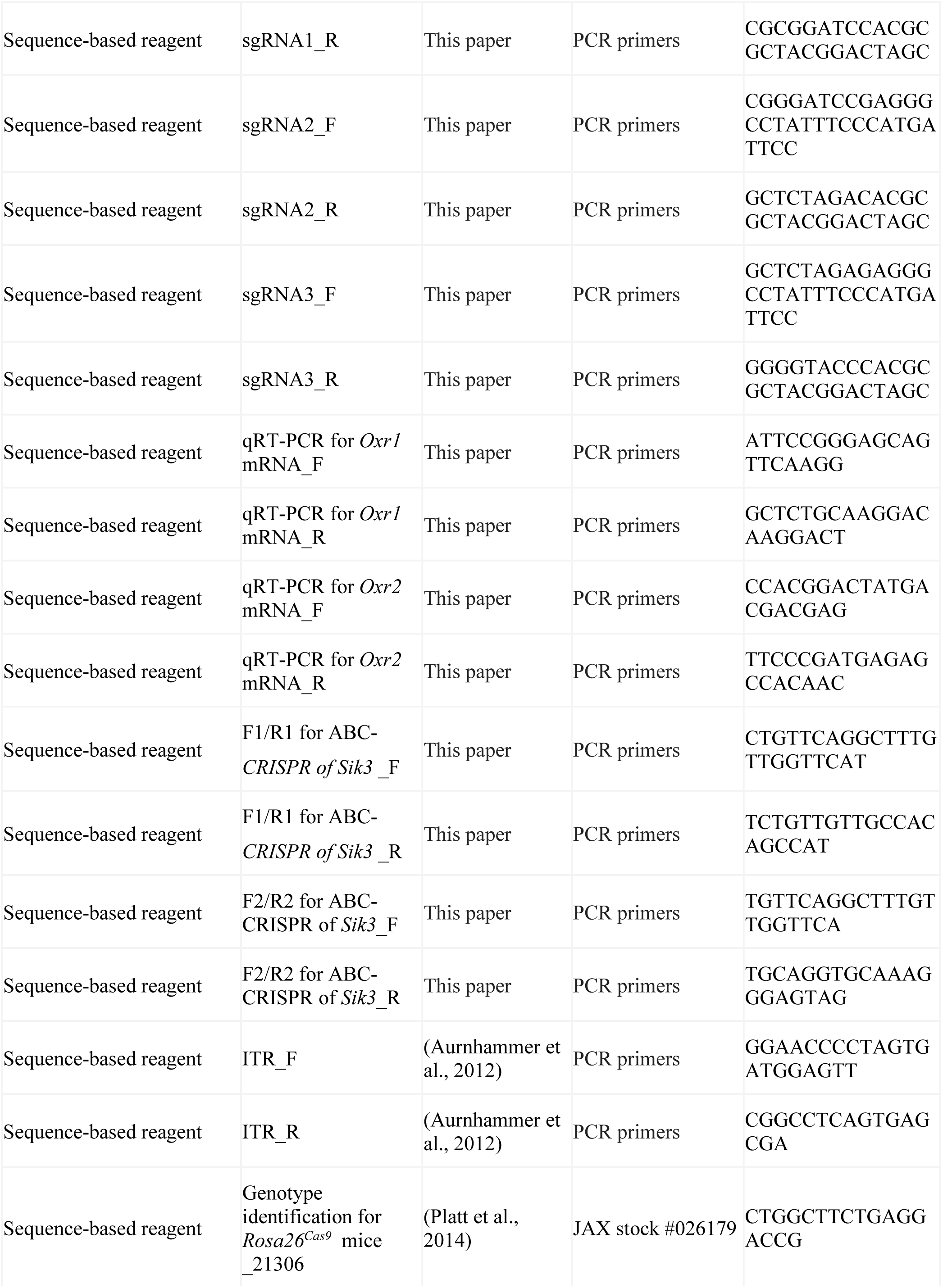
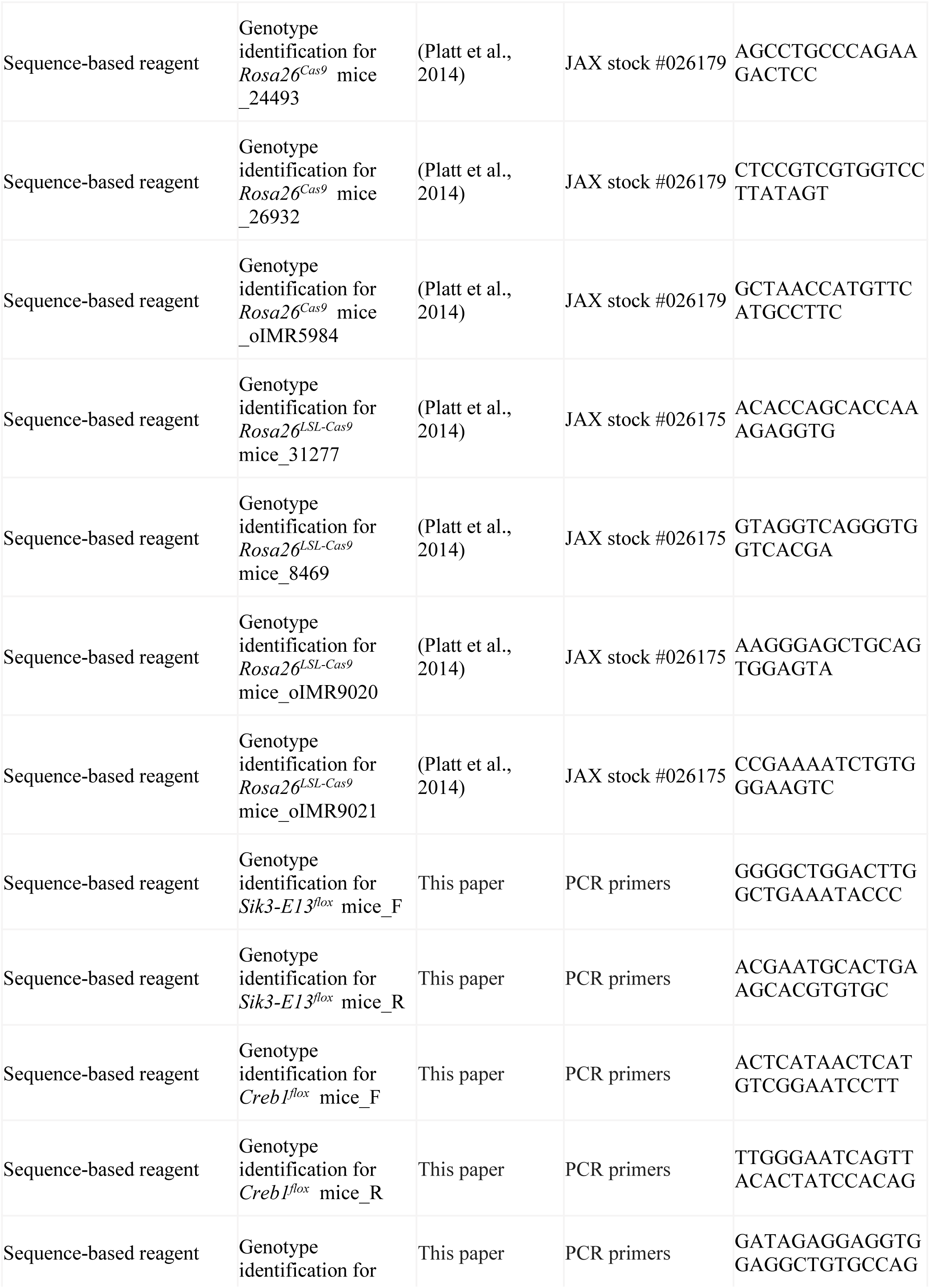
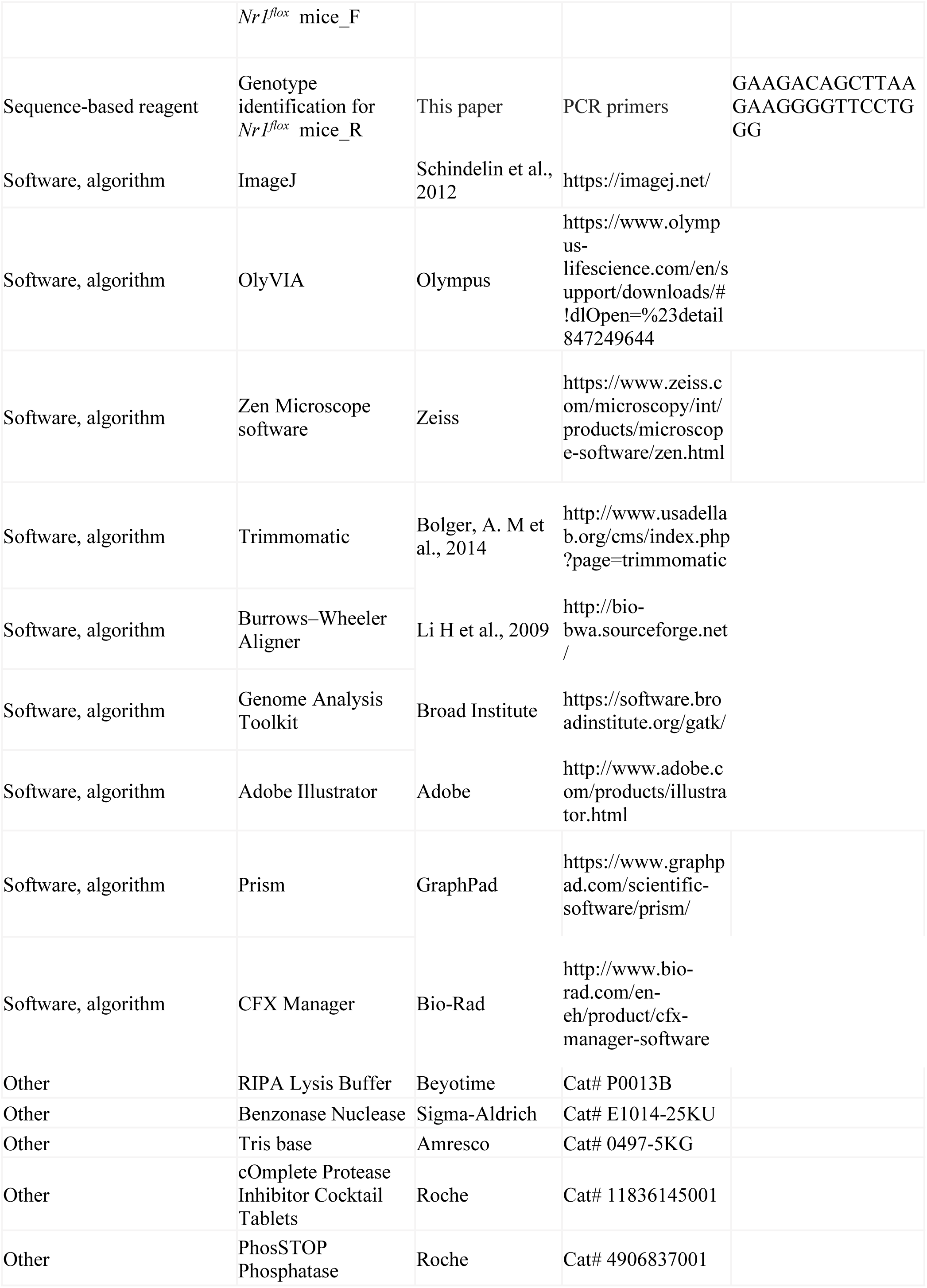
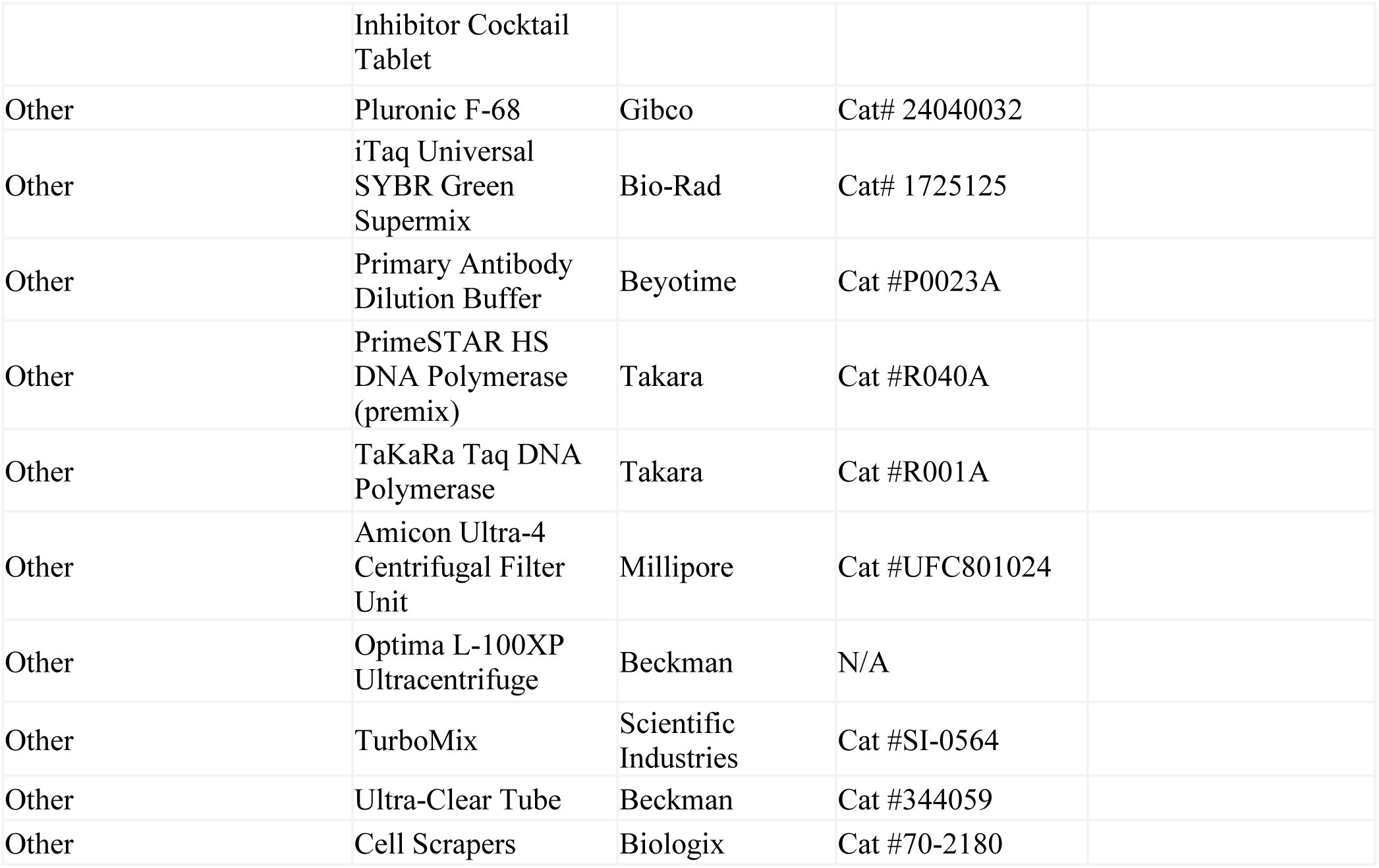

### Mice

All animal experiments were performed according to procedures approved by the institutional Animal Care and Use Committee of National Institute of Biological Sciences, Beijing (NIBS). All mice were provided food and water *ad libitum* and were housed under humidity and temperature controlled conditions (22-24℃) on a 12-h light: 12-h dark cycle. *Rosa26^Cas9^* (JAX, 026179) and *Rosa26^LSL-Cas9^* (JAX, 026175) were purchased from the Jackson laboratory. The *Creb1^flox^*, *Sik3-E13^flox^* and *Nr1^flox^* mice were generated by flanking exon 4 of *creb1*, exon13 of *Sik3* or exons 7 and 8 of *Nr1* with two loxP sites, respectively, in the transgenic animal facility at NIBS. All strains were bred to generate the appropriate genotypes for this study. Littermates were used wherever possible for experiments. Most experiments used male mice except *Sik3-E13^flox^;Rosa26^Cas9^* female mice which were specific indicated in the study. For each experiment, the age of the animals is ranged from 11 to 18 weeks.

### Generation of AAV vectors

The pAAV: EF1α-DIO-H2B-eGFP plasmid was a gift from Dr. Minmin Luo’s lab (NIBS). pAAV-CBh-Cre was constructed by cloning the CBh-Cre cassette from pAAV-U6-sgRNA(backbone)-CBh-Cre (Addgene, 60229) into MluI-HF (R3198L, NEB) and HindIII-HF (R3104S, NEB) digested pAAV-hSyn-eGFP (Addgene, 105539). pAAV-hSyn-Cre and pAAV-hSyn-SNIPPs were subcloned with AsiSI (R0630L, NEB) and MluI-HF (R3198L, NEB) restricted endonuclease from pAAV-hSyn-eGFP by replacing eGFP with Cre or Abi1 (Origene, MR206165), Anks1b (Origene, MR219888), Arfgap2 (Origene, MR208335), Arfgap3 (Origene, MR208404), Ankrd63 (Origene, MR213874), Synpo (Origene, MR215248), Ddx3y (Origene, MR209824), Pdha1 (Origene, MR206079) coding sequence.

The cDNAs of Arc (CCDS37102.1), Homer1a (CCDS26687.1), Rab3a (CCDS22379.1), Plk2-dn (CCDS26767.1, 332-682aa) , CamKIIα (CCDS29276.1), CamKIIβ (CCDS24411.1), CaMKIV (CCDS29122.1), CDK5 (CCDS51434.1), CREB (CCDS15005.1) and CRTC1 (CCDS40372.1) were amplified from mouse brain cDNA library by primers contain AgeI-HF (R3552L, NEB) and BamHI-HF (R3136L, NEB) digested site. CaMKIV-dn and CRTC1^CA^ were constructed from CaMKIV or CRTC1 by site-directed mutagenesis using specific primers that contain K75E or S151A/S245A mutations. MEF2^VP16^ and CREB^VP16^ were constructed by fusing MEF2C (CCDS84042.1, 1-117aa) and CREB (CCDS15004.1, 88-341aa) with C-terminal VP16-3xHA and N-terminal VP16 by Gibson assembly. All these cDNAs were inserted into pAAV-hSyn-eGFP by replacing eGFP with individual cDNA.

The pAAV-Tre-eGFP, pAAV-Tre-CREB^VP16^ and pAAV-Tre-CRTC1^CA^ were constructed by subcloing Tre promoter to pAAV-hSyn-eGFP, pAAV-hSyn-CREB^VP16^ and pAAV-hSyn-CRTC1^CA^. Reverse tetracycline-controlled transactivator (rtTA) expression was driven by the EF1α promoter. Tre promoter and rtTA sequences were obtained from Dr. Feng Shao’s lab (NIBS).

The sgRNA sequences were designed based on the Mouse GeCKO v2 Library (Sanjana et al., 2014; Sunagawa et al., 2016). Annealed oligos for individual sgRNA were first inserted into the SapI (R0569S, NEB) digested pAAV-U6-sgRNA(backbone)-hSyn-Cre-2A-EGFP-KASH (Addgene, 60231) vector. To better construct the double and triple sgRNA vectors, we had inserted an multiple cloning site (MCS) sequence before the hSyn promoter of pAAV-hSyn-eGFP and pAAV-hSyn-Cre. Then, these modified pAAV-hSyn-eGFP or pAAV-hSyn-Cre plasmids contain double or triple U6:sgRNA cistrons were constructed through PCR amplification two or three different sgRNAs that target the same gene by two or three pair primers, Restriction endonuclease of MluI-HF (R3198L, NEB), BamHI-HF (R3136L, NEB), XbaI (R0145S, NEB) and KpnI-HF (R3142S, NEB) were used for the vectors and fragments digestion, Quick Ligation Kit (M2200L, NEB) was used for the ligation of these appropriate clone. All these plasmids were saved and amplified in NEB Stable Competent *E. coli* (C3040I, NEB).

### Virus generation

AAV-PHP.eBs were packaged in AAVpro 293T cells (Clontech, 632273). Cells were harvested by cell lifter (Biologix, 70-2180) 72-h after co-transfection with PHP.eB (Addgene, 103005), pAdDeltaF6 (Addgene, 112867) and transfer plasmids using polyethylenimine MAX (Polysciences, 24765). The cell pellets were suspended in 1X Gradient Buffer (10mM Tris-HCl pH=7.6, 150mM NaCl, 10mM MgCl_2_). Cells were lysed by five repeated cycles of liquid nitrogen freezing, 37℃ water bath thawing and vortex. Then the cell lysate was mixed with ≥ 50 U/mL of Benzonase nuclease (Milipore, E1014) and incubated at 37℃ for 30 min. Centrifuge the cell lysate at 21,130g for 30 min at 4℃ and transfer the supernatant to a pre-build iodixanol (Optiprep, D1556) step gradients (15%, 25%, 40% and 58%) for ultracentrifugation purification. Vacuum centrifuge at 41000rpm, 4℃ for 4-h, the virus particles were in the layer of 40% iodixanol gradient. Accurately insert the needle ∼1-2 mm below the interface between the 40% and 58% gradient and extract all the 40% virus containing layer. Purified AAV-PHP.eB were concentrated using Amicon filters (EMD, UFC801096) and formulated in sterile phosphate-buffered saline (PBS) supplemented with 0.01% Pluronic F68 (Gibco, 24040032). Virus titers were determined by qPCR using a linearized AAV plasmid as a standard.

### Video-based sleep recording and screening

For video-based sleep screening, 9-11 weeks old C57BL/6J male mice were retro-orbitally injected with 10^12^ AAV-PHP.eB viruses expressing different cDNAs from the hSyn promoter. Two weeks after virus injection, mice were individually housed in Ancare cages with food and water provided *ad libitum* on a 12-h light: 12-h dark cycle. The sleep/wake behaviors of the mice were recorded by an infrared camera (704 x 576 resolution) at 25 frames/s. Infrared LED lights were placed above the cages for videotaping mouse behaviors during the dark phase. For all mice, 3 days of video data were continuously recorded to calculate average daily sleep time for each mouse.

### Automatic sleep/wake staging by SleepV software

The automatic sleep/wake staging of video data consists of two stages: 1) an AI-augmented video analysis algorithm was developed to extract various information of test mouse from each sampled frame. 2) the state of test mouse (active or inactive) at the video clip level was obtained by grouping the frame-level information within a time window.

#### (1) Video analysis stage

The suspected regions of interest (ROIs) are first extracted from each sample frame using traditional image processing techniques, such as Gaussian filtering, adaptive and global threshold. More than one ROIs are allowed to be detected because of the complication of food, water gel and shadows caused by environmental light. This multi-detection strategy ensures that the true mouse region is not missed. A deep neural network is then used to make a binary classification of each ROI as a mouse or not. Our neural network backbone was based on Resnet18 architecture (He et al., 2016). We trained the Resnet18 model using 7,200 positive images and 9,000 negative images with manual labeling. All image frames for training were randomly chosen from six mice, which covered both light and dark conditions, and as many mouse postures as possible. All the experiments were carried out on a workstation with a NVIDIA Titan XP GPU, Intel Core i7-7000 CPU.

During the training process, labeled frames data were randomly split into two parts: 80% for training and 20% for validation. We chose cross entropy as the binary classification loss function, and Stochastic Gradient Descent (SGD) as the optimizer with a fixed learning rate 0.001. The training process lasted 50 epochs, as a result, the model accuracy was 99.3% on training dataset and about 96.1% on validation dataset. The ROI with the highest confidence score is designated as the mouse region, while the others were classified as the background noise.

According to the size of ROI and position coordinate, we extracted several descriptive features, i.e., the network prediction score (predict), the detected mouse mask area (IoU), the center of mass of the mask (activity), and the gray information (color) within the detected mask. Similarly, the frame-level features within a time window centered on that frame were also extracted. Denote f_{t} as the current frame, {f{t-m}, f{t-m+1}, … f{t+m}} refer to the frames within this time window, where m is a hyperparameter. The differences between f_{t} and f{t+i}, including predict score difference P_i (predict), mask difference M_i (IoU), center of mass difference of the mouse body A_i (activity), and gray pixel difference G_i (color), were calculated and normalized. We defined P_i >0.1, or G_i >3, or A_i >5, or the normalized value (G_i + M_i + P_i)/3>0.5 as the threshold to judge the mouse was active at frame t.

#### (2) Sleep/wake state determination stage

According to the frame-level preliminary judgment, SleepV defines the sleep state as ≥ 40-s of continuous immobility as previously developed video-based sleep analysis softwares (Fisher et al., 2012; Pack et al., 2007). To further improve the performance of SleepV, we used a series of 0, 5, 10, 15, 20, 25, 30s thresholds to filter subtle mouse movements during sleep and compared the sensitivity, specificity, accuracy, and sleep bouts number of SleepV with EEG/EMG analysis. We found that annotating ≤ 15-s of mouse movements in between two sleep states as sleep achieve the best performance of automatic sleep/wake staging by SleepV, which is comparable to that of semi-automatic sleep/wake staging by EEG/EMG analysis software with manual corrections.

### Tet-on inducible system

For Tet-on inducible sleep recording, EEG/EMG surgeries were carried out on 12-weeks old mice. The mice were recovered for one week and AAV-Tre-eGFP, AAV-Tre-CREB^VP16^ or AAV-Tre-CRTC1^CA^ was co-injected with AAV-EF1α-rtTA through retro-orbital injection. After two weeks virus expression, baseline sleep recording was performed for 3 days without Doxycycline. Then the water was changed with 2mg/mL Doxycycline containing water and subjected to EEG/EMG recording for another 3 days.

### Genomic DNA Extraction and Captured Illumina Sequencing

Genomic DNA was extracted from mouse brain using TIANamp Genomic DNA Kit (TIANGEN, DP304) following the recommended protocol. Genomic DNA (1-1.5 g) was sheared to 300 to 400-bp by Covaris S220 (Covaris, Woburn, MA, USA) and purified with 1X magnetic beads (Ampure XP; Beckman Coulter). Sheared DNA fragments was Subjected to Illumina paired-end DNA library using NEBNext ultra II DNA Library Prep Kit (E7645L, NEB). Preparation and PCR-amplified for three cycles and libraries size were selected with 0.55-1X magnetic beads (Ampure XP, Beckman). Amplified libraries were sequenced using the HiSeq X ten Platform (Illumina) as paired-end 150 base reads according to the manufacturer’s protocol.

Illumina raw sequencing reads were processed through a standard pipeline consisting of low-quality read filtering through Trimmomatic (version 0.36), alignment to mouse genome GRCm38 (mm10) using the Burrows–Wheeler Aligner (BWA, version 0.7.17-r1188) algorithm. The aligned BAM files were processed using the Genome Analysis Toolkit (GATK, version 4.1.4), including mark PCR duplicates and correction for realignments and mapping quality score recalibrations. Haplotype Caller was used for variant calling.

Twenty-one potential off-target sites for the three sgRNAs targeting *Sik3* and *NeuN* genes listed below were identified using Cas-OFFinder (http://www.rgenome.net/cas-offinder/).

Sik3 sgRNA-1(ACAGACCACTTGGTAGCCCA):

1. AagGACCACTTGtTAGCCCAAGG chr2:87098654

2. ACAaAgCACTTGGTAGCCtAAGG chr4:46756223

3. ACAGACCACgTGtTAGaCCAAGG chr4:57954753

4. gCAGAaCACTGTGGTAGCCCAGGG chr4:126131919

5. AtA-ACCACTTGGTAGgCCAGGGAGG chr17:86795878

6. ACAGACCAaTctGT-GCCCAGGGTGG chr8:122648253

7. ACAGACCACaTGG-AGCtCAAGaAGG chr15:5662796

Sik3 sgRNA-2(ACTCACCCATATGTCCACTT):

8. ACTCAgCCATtTGTCCAtTTGGG chr7:123562465

9. ACTCACtCATATGTgCACTgGGG chr2:25603475

10. ACTCACCCtTAgGgCCACTTGGG chr12:17052697

11. ACaCAgCCAgATGTCCACTTGGG chr17:60959507

12. ACTCACaaATATG-CCAgTTTGGTGG chr15:16585273

13. tCTCAgCCATATGTCCACTTC-cAGG chr5:136277718

14. A-TCACCCtTATGTgCtCTTTGGTGG chr7:66324551

Sik3 sgRNA-3(GTTCAAACAGATCGTCACAG):

15. GTTCAAACAGATCGaaACgGAGG chr5:52797383

16. GTTCAcACAGtTaGTCACAGTGG chr5:144210416

17. GTTCtAACAaATtGTCACAGGGG chr1:49457652

18. GcTCAAACAtATtGTCACAGGGG chr4:17192911

19. GTCTCAAACAcATCGgCACAGGGG chr8:4772824

20. GTCTCAAACAcATCGaCACAGGGG chr1:163033701

21. GaTCAAACAGATaGT-ACAGAGG chr15:77931180

NeuN sgRNA-1(TCGGGGTCCCTGAACCGGAA):

1. TCtGGaTCCCgGAACCGGAAAGG chr16:7277126

2. TCGGGGTCCCTGAACCactAAGG chr7:30375662

3. TaGGGGTCCCTGAAaCaGAATGG chr7:45640152

4. TgGGGtTCCCTGAACCccAAAGG chrX:11971373

5. TCTGGGGTCCCTGAACCtGcAAGG chr17:48361361

6. TCGGGGTCCCaGCAACCaGAAGGG chr5:99874083

7. TCGGGGaCCCgGAACCTGGAAGGG chr5:109528520

NeuN sgRNA-2(GCTCAGATGCTGACCGAGCC):

8. GCTCAGATGCTGACaGAcCtGGG chr16:25376832

9. GCTgAGATGCaGACtGAGCCTGG chr16:30614345

10. GCTCAGcTGCTGgCCcAGCCTGG chr15:82930003

11. GCTtAGATGCTGAtgGAGCCTGG chr7:101383685

12. GCTCAGATGCTGgCCTGtGCCAGG chr9:120120623

13. GCTCAGATGtTGtCCAGAGCCAGG chr5:66096471

14. GCTCAGGcTGCTGACaGAGCtGGG chrX:11232414

NeuN sgRNA-3(GCTGAATGGGACGATCGTAG):

15. GCTGttTGGGAgGATCGTAGAGG chr13:48330995

16. GCTGtATGGtAgGATCGTAGGGG chr4:22110214

17. GCTGAATGGGAgGATCaTcaGGG chrX:98480354

18. GCTGAATGGagCGATgGgAGAGG chr15:10917958

19. GCTGGAATGtGAgGATaGTAGTGG chrX:164503183

20. GCTGAATGAGGAaGATaGTgGTGG chr5:35422881

21. GCTGgATGGGACGATgGTACtTGG chr8:28174078

### EEG/EMG surgery

All EEG/EMG surgeries were performed by experienced technicians. 11 to 13-weeks old male or female mice (all experiments were conducted with male mice if not indicated otherwise) were anesthetized by isoflurane (4% for induction, 2% for maintenance) and surgical tools were sterilized by ethanol just before use. After confirming the mice lack of pain, the head region was shaved, cleaned with ethanol and the skull was exposed. The exposed skull was cleaned by cotton swabs to improve binding of skull and dental cement. Handheld electrical drill was moved to the lambda point and set the coordinate as (0, 0, 0). Then four holes were drilled by the electrical drill in the skull. The coordinate of the holes were (-1.27, 0, 0), (-1.27, 5.03, 0), (1.27, 5.03, 0) and (1.27, 0, 0). Then the EEG electrode pins were implanted to the dura under stereotaxic control and the EMG wires were inserted into the neck muscle and then stick to the skull with dental cement. After surgery, the mice were housed individually to recover for one week. Then retro-orbital injection of AAV-PHP.eB was performed. After allowing time for virus expression (2 week for ABC expression and 3 weeks for ABC-KO), the mice were tethered to a counterbalanced arm (Instech Laboratories) that allowed free movement and exerted minimal weight for one week before EEG/EMG recording.

### EEG/EMG recording and data analysis

Three days of baseline EEG/EMG recording were conducted after mice were acclimated for one week recording condition. For the ABC-*Ox1r/Ox2r^DKO^* mice, an additional one week EEG/EMG recording were conducted feeding with chocolates. The sleep–wake behaviors were analyzed as previously described with modifications (Funato et al., 2016). EEG/EMG data were visualized and analyzed using a custom semi-automated staging MatLab (MathWorks)-based program, followed by visual inspection. Following semi-automated analysis of EEG/EMG data, EEG signals were subjected to fast Fourier transform analysis for 1 to 30Hz with 1-Hz bins. Sleep/wake state was staged into NREMS, REMS and wake. NREMS was staged by high amplitude, delta (1–4 Hz) frequency EEG and low EMG tonus. REMS was characterized based on theta (6–9Hz)-dominant EEG and EMG muscle atonia. Wake was staged based on the presence of low amplitude, fast EEG and high amplitude, variable EMG signal. For the NREMS delta power density analysis, hourly averages of delta density were defined by the ratio of delta power (1–4 Hz) to total power of NREMS EEG. For the power spectrum analysis of NREMS, REMS and wake, the EEG power of each frequency bins was expressed as the percentage of total EEG power over all frequency bins (1–30 Hz) in each state.

### Brain lysate preparation and Immunoblotting

Mouse brains were quickly dissected and flash frozen in liquid nitrogen. Brain tissues were homogenized using mortar/pestle with liquid nitrogen and then lysed in ice-cold RIPA buffer (50mM Tris-HCL pH=7.4, 150mM NaCl, 1% Triton X-100, 0.1% SDS) (Beyotime, P0013B) freshly supplemented with protease inhibitor (Roche, 11836145001) and phosphatase inhibitor (Roche, 4906837001) for 30 min and centrifuged at 21,130g for 15 min at 4℃. The supernatant was transferred to a new tube and boiled at 95℃ for 10 min with SDS-loading buffer. Western blotting was performed according to standard protocols using the following antibodies. Rabbit polyclonal anti-SIK3 antibodies were generated using Abcam custom antibody production service. The following antibodies were purchased from commercial sources: anti-CREB (CST, 9197S), anti-NeuN (Milipore, ABN78), anti-β-ACTIN (Beyotime, AF003), anti-Tublin J (CST, 5568), anti-GFP (Beyotime, AG279), anti-Cas9 (Abcam, ab204448), anti-HA (Sigma, H6533), anti-NR1 (Millipore, 05-432), anti-GAPDH (EarthOx, E021010-01), HRP-labeled Goat Anti-Rabbit IgG(H+L) (Beyotime, A0208), HRP-labeled Goat Anti-Mouse IgG(H+L) (Beyotime, A0216), AP-labeled Goat Anti-Rabbit IgG (H+L) (Beyotime, A0239), AP-labeled Goat Anti-Mouse IgG(H+L) (Beyotime, A0258). BeyoECL Plus (Beyotime, P0018M) was used for the coloration of Horseradish Peroxidase-based secondary antibodies. BCIP/NBT Alkaline Phosphatase Color Development Kit (Beyotime, C3206) was used for the coloration of Alkaline phosphatase-based secondary antibodies.

### Immunohistochemistry

Mice were deeply anesthetized by choral hydrate and perfused transcardially with 0.9% normal saline followed by 4% paraformaldehyde in PBS. Brains were post-fixed in 4% paraformaldehyde in PBS at room temperature for at least 4 hours followed by incubation in 30% sucrose in PBS at room temperature for 24 h. The cryo-protected brains were sectioned at 40 micron on a cryostat microtome (Leica). After washing in PBST(0.3% Triton X-100 in PBS) for 5 min three times, brain sections were incubated in blocking solution (3% BSA, 0.3% Triton X-100 in PBS) at room temperature for 1-h. Then brain sections were incubated with the primary antibodies overnight at 4 ℃ and immunofluorescence tagged secondary antibodies at room temperature for 2-h. After staining, the brain sections were mounted on adhesion microscope slides (Genview) and encapsulated in sealed tablets containing 3 mg/mL DAPI (Solarbio, C0060).The following antibodies were used: anti-HA (1:500, 11867423001, Roche), anti-NeuN (1:500, ABN78, Milipore), anti-GFP (1:2000, ab13970, Abcam), Alexa Fluor 555 conjugated goat-anti-rat (1:500, 4417, CST), Alexa Fluor 488 conjugated goat-anti-rabbit (1:500, 111-545-003, Jackson Immunoresearch), Alexa Fluor 488 conjugated goat-anti-chicken (1:500, ab150169, Abcam).

### RNA extraction and RT-PCR

Brain tissues from OX1R/OX2R double knockout mice and control mice were quickly collected in liquid nitrogen. Previously sterilized mortar and pestle were used to pulverize about the whole brain sample in liquid nitrogen. Until a fine powder was obtained, the samples were quickly transferred to a new cold 1.5-mL centrifuge tube. Then, 1000 µL TRIzol (Invitrogen, 15596026) was added and mixed by vortexing the tube to help cell lysis from tissue samples. Centrifuge 5 minutes, 12,000 x g at 4 ℃ to remove high fat content. After incubating 5 minutes at 4 ℃, 200 µL, chloroform was added to the samples and flipped centrifuge tube to mixing reagents. Incubate 3 minutes and then Centrifuge the samples for 15 minutes at 12,000 x g at 4 ℃. Transfer the Upper aqueous phase to a new tube and adding 500 µL isopropanol. Flip centrifuge tube to mixing reagents and incubating 10 minutes. Centrifuge the samples for 10 minutes at 12,000 x g at 4 ℃ and discarding the supernatant. Resuspend the pellet in 1000 µL 75% ethanol and vortex briefly, then Centrifuge for 5 minutes at 7500 x g at 4 ℃. Discard the supernatant and air dry the RNA pellet for 10 minutes. Resuspend the pellet with 200 uL RNase-free water. The RNA concentration of these samples were quantificated by NanoDrop One (Thermo Scientific, ND-ONE-W). 2 ug RNA per sample were used to reverse transcript the cDNA by QuantScript RT Kit (TIANGEN Biotech, KR103). 2 x iTaq Universal SYBR Green Supermix (Bio-Rad, 1725125) was used for the Reverse Transcription-Polymerase Chain Reaction (RT-PCR) by CFX96 Touch qPCR Detection System (Bio-Rad, 1855196). Data was analysed by CFX Manager software (Bio-Rad).

### Quantification and statistical analysis

Statistical analysis of EEG/EMG data was performed using GraphPad Prism 8.0.2. Knockout efficiency measured by Western blot was quantified using ImageJ software. Two-tailed P value was used for unpaired t-test, one-way ANOVA for multiple comparisons and two-way ANOVA for multiple comparisons involving two independent variables. Dunnett’s test compares every mean to a control mean. *p* < 0.05 was considered statistically significant.

## Notes

https://bigd.big.ac.cn/gsa/browse/CRA005128

https://github.com/wochiguodong/SleepV.git

### Competing Interest Statement

The authors have declared no competing interest.

